# NCoR1 and SMRT fine-tune inflammatory versus tolerogenic balance in dendritic cells by differentially regulating STAT3 signaling

**DOI:** 10.1101/2021.03.11.434976

**Authors:** Atimukta Jha, Abdul Ahad, Gyan Prakash Mishra, Kaushik Sen, Shuchi Smita, Aliva P Minz, Viplov Kumar Biswas, Archana Tripathy, Shanti Bhushan Senapati, Bhawna Gupta, Hans Acha Orbea, Sunil Kumar Raghav

**Author notes:** Equal contribution first author (Atimukta Jha, Abdul Ahad, Gyan Prakash Mishra). Corresponding author: Sunil Kumar Raghav, PhD., Email for correspondence.

## Abstract

Dendritic cell (DC) fine-tunes inflammatory versus tolerogenic responses to protect from immune-pathology. However, the role of co-regulators in maintaining this balance is unexplored. NCoR1-mediated repression of DC immune-tolerance has been recently reported. Here we found that depletion of NCoR1 paralog SMRT (NCoR2) enhanced cDC1 activation and expression of IL-6, IL-12 and IL-23 while concomitantly decreasing IL-10 expression/secretion. Consequently, co-cultured CD4^+^ and CD8^+^ T-cells depicted enhanced Th1/Th17 frequency and cytotoxicity, respectively. Comparative genomic and transcriptomic analysis demonstrated differential regulation of IL-10 by SMRT and NCoR1. SMRT depletion represses mTOR-STAT3-IL10 signaling in cDC1 by downregulating NR4A1. Besides, *Nfkbia* and *Socs3* were down-regulated in *Ncor2* (*Smrt*) knockdown cDC1, supporting increased production of inflammatory cytokines. Moreover, studies in mice showed, adoptive transfer of SMRT knockdown cDC1 in OVA-DTH induced footpad inflammation led to increased Th1/Th17 and reduced tumor burden after B16 melanoma injection by enhancing oncolytic CD8^+^ T-cell frequency, respectively. We also depicted decreased *Ncor2* expression in Rheumatoid Arthritis, a Th1/Th17 disease.

## Introduction

Dendritic cells (DCs) play an important role in immune surveillance and maintain an optimal balance between inflammation and immune-tolerance to avoid immune-pathology [1,2]. After encountering pathogens DCs undergo activation and maturation leading to secretion of cytokines and expression of co-stimulatory molecules, which ultimately decides the fate of a particular T-cell response to clear the invading pathogens [3]. Hence, antigen specific activation of DCs is an important event to orchestrate the immune system culminating into development of T-cell adaptive response for the induction and expansion of either an optimum protective pro-inflammatory or tolerogenic response [4]. DC subtype specific differential expression of toll like receptors (TLRs) along with their unique signaling cascades provide the specificity against the pathogens e.g., plasmacytoid DCs (pDCs) highly express TLR7 and TLR9, required to mount antiviral response whereas conventional DCs (cDCs) are mainly responsible to maintain the balance between inflammatory vs tolerogenic response and cross presentation [5–7]. Among cDCs, cDC1s (CD8α^+^ DCs) are considered as bystanders which integrate signals derived from intracellular infection to tailor the appropriate CD4^+^ T-cell response along with anti-tumor CD8^+^ T-cell activation with the help of their unique property of cross presentation [8]. cDC1s identify pathogen associated molecular patterns (PAMPS), such as bacterial unmethylated CpG-DNA or viral double stranded RNA through TLR9 and TLR3 respectively [9]. Upon TLR9 stimulation in cDC1s, the TIR domain of TLR and an adapter, Myd88, activates interleukin-1 receptor-associated kinase-4 (IRAK4) and IRAK1 [10]. IRAK4 subsequently activates the NF-kB signaling [11]. The TLR9-Myd88 signaling has also been linked with JAK-STAT signaling pathway for cytokines production [12]. STAT3 is a new entry in the block although the TLR-Myd88-STAT3 signaling does not affect NF-kB signaling in B-cells [13]. STAT3 depleted DCs have been shown to be insensitive to IL-10 mediated suppression leading to hyper-activation of T-cells and inflammation in mice [14]. A combination of such divergent DC signals leads to differentiation of naive T helper cells to various effector subtypes such as Th1, Th2, Th17 or Tregs. A fine balance of secretory cytokines like IL-6, IL-12, and IL-23 modulates Th cells towards Th1 or Th17 subtypes, whereas increased levels of IL-10, SOCS3 and CD83 differentiate them towards Tregs [15, 16]. Thus, a tight regulation of these cytokines is important to maintain the fine balance between inflammatory, anti-inflammatory or tolerogenic responses [17]. The idea of perturbing T-cell differentiation to modulate the immune system for cell mediated therapy is an accepted concept. For example, the use of DCs to manipulate T-cells in cancer immunotherapy has been widely reported. Despite a number of attempts, there are multiple occurrences of insufficient T-cell activation and effective priming in *in vivo* systems [8]. Therefore, identifying ways to perturb DC responses in a controlled manner to enhance T-cell function is an interesting area. Recently, NF-kB signaling was shown to be perturbed in tolerogenic DCs having increased IL-10 production [18]. However, the regulators underlying the fine modulation of these TFs are not clearly documented. A group of TFs belonging to nuclear receptors (NRs) such as NURR-77 or NR4A1 and PPAR-γ are reported to have a role in regulation of cytokine gene expression [19]. NURR-77 controls production of IL-6, TNF-α, and IL-12 in both human and murine dendritic cells [20]. Similarly, peroxisome proliferator-activated receptor (PPAR-γ) also exerts anti-inflammatory effects in monocytes and macrophages [20]. Although TFs and NRs lead to activation of transcription, their activity is tightly regulated by a network of co-regulators, including co-activators and corepressors. For example, co-regulators ASC-2 and SMRT control activation and repression of Nur77 respectively [21]. While ASC-2 is dependent on CaMKIV for the transactivation of Nur77, SMRT on the contrary binds directly to Nur77, through its C-terminal domain nuclear receptor interaction motif, and represses it [21]. All these observations are made in HeLa and CV-1 cells. Nuclear receptor co-repressor 1 (NCoR1) and its paralog Silencing mediator of retinoic acid and thyroid hormone receptor (SMRT) were identified in relation to unliganded thyroid and retinoic acid receptor mediated repression of gene expression [22]. *Ghisletti et. al.* has shown that the combination of NCoR1 and SMRT is required for regulation of inflammatory and anti-inflammatory genes in macrophages [23]. Recent report showed that siRNA mediated depletion of SMRT in IL4 and GM-CSF differentiated DCs led to decreased expression of CD209 mRNA [24]. However, the combinatorial role of NCoR1 and SMRT in immune response regulation in DCs is largely unexplored. Recently we demonstrated that active repression of tolerogenic genes like IL-10 by NCoR1 is essential for development of immunogenic response in DCs [25]. NCoR1 depletion enhanced the expression of tolerogenic molecules like IL-10, IL-27, PDL1 and SOCS3 resulting in increased frequency of Tregs and shift of immunogenic balance towards tolerance [25].

In this study, we explored the role of SMRT in modulating the immune function of cDC1 DCs *in vitro, ex vivo* and *in vivo*. We identified how two highly homologous nuclear receptor co-repressor proteins NCoR1 and SMRT tightly control the fine balance of inflammatory and tolerogenic response in DCs, which consequently regulates the differentiation of naïve Th cells into Th1, Th17 or Tregs. Moreover, comparative genomewide binding of NCoR1 and SMRT and transcriptomic analysis of NCoR1 and SMRT depleted cDC1 revealed their differential role in control of STAT3 signaling and IL-10 expression and its underlying control of repression. Overall, our study demonstrated that NCoR1 and SMRT are potential targets for regulating a fine balance of DC mediated inflammatory and tolerogenic T-cell responses.

## Results

### SMRT KD cDC1 DCs depicted enhanced activation / co-stimulation

We observed that Nuclear Receptor Co-repressor-2 (*Ncor2*) was constitutively expressed at transcript levels in murine cDC1 (mutu-cDC1 line) before and after 2h, 6h and 12h of activation by TLR9 ligand, CpG-B [26] (**Supplementary Figure 1A**). To identify the potential role of SMRT in cDC1, we generated stable *Ncor2* gene knock down (SMRT KD) and empty vector transduced (control) cDC1 mutu-cDC1 lines. *Ncor2* gene depletion was confirmed at transcript (>75-80%) and protein level (30-50%) in unstimulated and (~75-80%) in 6h CpG challenged conditions (**Figure 1A and Supplementary Figure 1B**). Then we performed RT-qPCR for pro-inflammatory and anti-inflammatory cytokine gene transcripts (*Il12b* & *Il10*) in control and SMRT KD cDC1 cells before and after 2h and 6h CpG challenge. We found significantly increased *Il12b* with a concomitantly decreased *Il10* post CpG stimulation in SMRT KD cDC1 compared to control cells (**Figure 1A**). Besides, flow cytometric analysis showed significantly increased percentage positive cells for CD80, CD86 and CD40 in SMRT KD cDC1 as compared to controls (**Figure 1B**). The median fluorescence intensity (MFI) shifts also showed a similar trend (**Supplementary Figure 1C**). We observed that SMRT KD cDC1 preserved the similar phenotype under TLR3 stimulation with pIC as well (**Supplementary Figure 1D**). Next, we investigated the antigen presentation ability by assessing expression of MHC-I and MHC-II on SMRT depleted DCs. We found significantly increased MHC-I percent positive cells in both CpG and pIC activation whereas MHC-II levels remained unchanged (**Figure 1B and Supplementary Figure 1D**). MFI shifts depicted similar trends. Gating strategy used for the mutuDC cell line is uniform for all experiments (**Supplementary Figure 1E**). To further validate the impact of SMRT depletion in primary *ex vivo* cDC1s, we performed transient SMRT depletion in primary bone-marrow derived cDC1s (BMcDC1s) cultured in FLT3L. We found that CD80 and CD86 showed an increased trend in SMRT KD condition as compared to controls (**Figure 1C and Supplementary Figure 2A**). However, we observed a non-significant decrease in MHC-II upon SMRT depletion in primary cDC1s (**Supplementary Figure 2A**). Uniform gating strategies were used throughout BMcDC1analysis (**Supplementary Figure 2B**).

**Figure 1.**
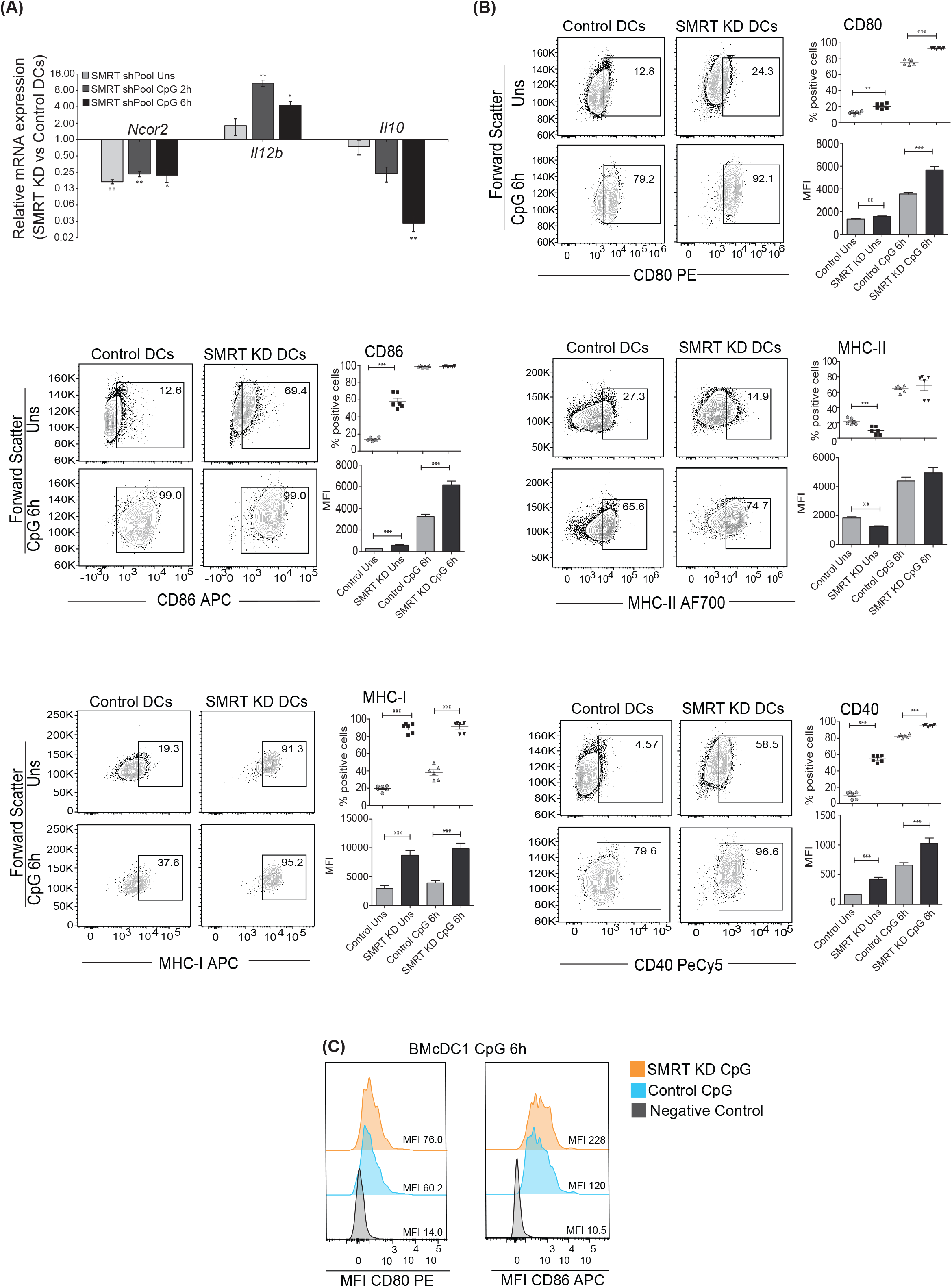
SMRT depleted cDC1 and BMcDC1 exhibit enhanced activation and maturation upon TLR9 ligation with CpG-B. **A.** RT-qPCR showing relative transcript expression of *Ncor2, Il12b* and *Il10* in unstimulated, 2h and 6h CpG-B stimulated control and SMRT KD cDC1. (n=3) **B.** Flow cytometry analysis of co-stimulatory surface markers CD80, CD86, MHC-II, MHC-I, and CD40 in unstimulated and 6h CpG stimulated control and SMRT KD cDC1 mutuDCs. Corresponding contour plots, dot plots, and bar-plot showed the percent positive cells and MFI shifts for each of the marker genes. (n=6) **C.** Flow cytometry analysis depicting MFI histograms of co-stimulatory molecules CD80 and CD86 in 6h CpG stimulated control and SMRT KD bone-marrow derived cDC1 (BMcDC1). (n=8) *p ≤ 0.05, **p ≤ 0.01 and ***p ≤0.001. p-value has been calculated using two tailed paired student’s t-test. Data shown in figure is combined from 3 independent experiments [A-D]. Error bars represent SEM.

### SMRT depleted cDC1 showed increased IL-6, IL-12 and IL-23 pro-inflammatory cytokines with concomitantly decreased IL-10

Next, we assessed the expression of important DC response cytokines in both, mutu-cDC1 line and primary BMcDC1s. We found that SMRT depletion significantly enhanced the percent positive cells as well as MFI shifts for IL-6, IL-12p40 and IL-23p19 cytokines after 6h CpG challenge as compared to control cDC1. IL-23p19 was significantly increased in SMRT KD cDC1 irrespective of stimulation (**Figure 2A and Supplementary Figure 3A**). On the contrary, we found that IL-10 was significantly reduced upon activation in SMRT KD DCs (**Figure 2A and Supplementary Figure 3A**). Similar results were observed after TLR3 stimulation with pIC (**Supplementary Figure 3B**). Uniform gating strategies were used throughout cDC1 analysis (**Supplementary Figure 3C**). Further to estimate the levels of these cytokines in CpG activated cDC1 culture supernatants, we performed bio-plex assays and found the cytokine levels of IL-6, IL-12p40 and IL-12p70 were significantly increased in SMRT depleted mutu-cDC1 line with a drastic reduction in IL-10 (**Figure 2B**). Moreover, the IL-2 cytokine necessary for clonal expansion of T-cells was found to be significantly increased in SMRT KD cDC1 compared to controls (**Figure 2B**). We further confirmed these findings in primary BMcDC1s and similar to our mutuDC line, we found significantly increased IL-6 and IL-23 and decreased IL-10 cytokine in BMcDC1s as well (**Figure 2C and Supplementary Figure 4A**). However, IL-12p40 percent positive cells showed a non-significant decrease after SMRT KD in BMcDC1s (**Supplementary Figure 4A, Supplementary Figure 4B**). Although IL-12p40 percent positive cells in BMcDC1s was reduced in KD compared to control however, similar to mutuDC line data, bio-plex cytokine assays in control and SMRT KD BMcDC1s showed a significant increase in secreted IL-12p40 after 6h CpG-B stimulation in KD cells (**Figure 2D**). IL-6, IL-12p70, and IL-2 also showed a nonsignificant but increasing trend (**Figure 2D**). These observations suggested a strong inflammatory phenotype for SMRT KD cDC1.

**Figure 2.**
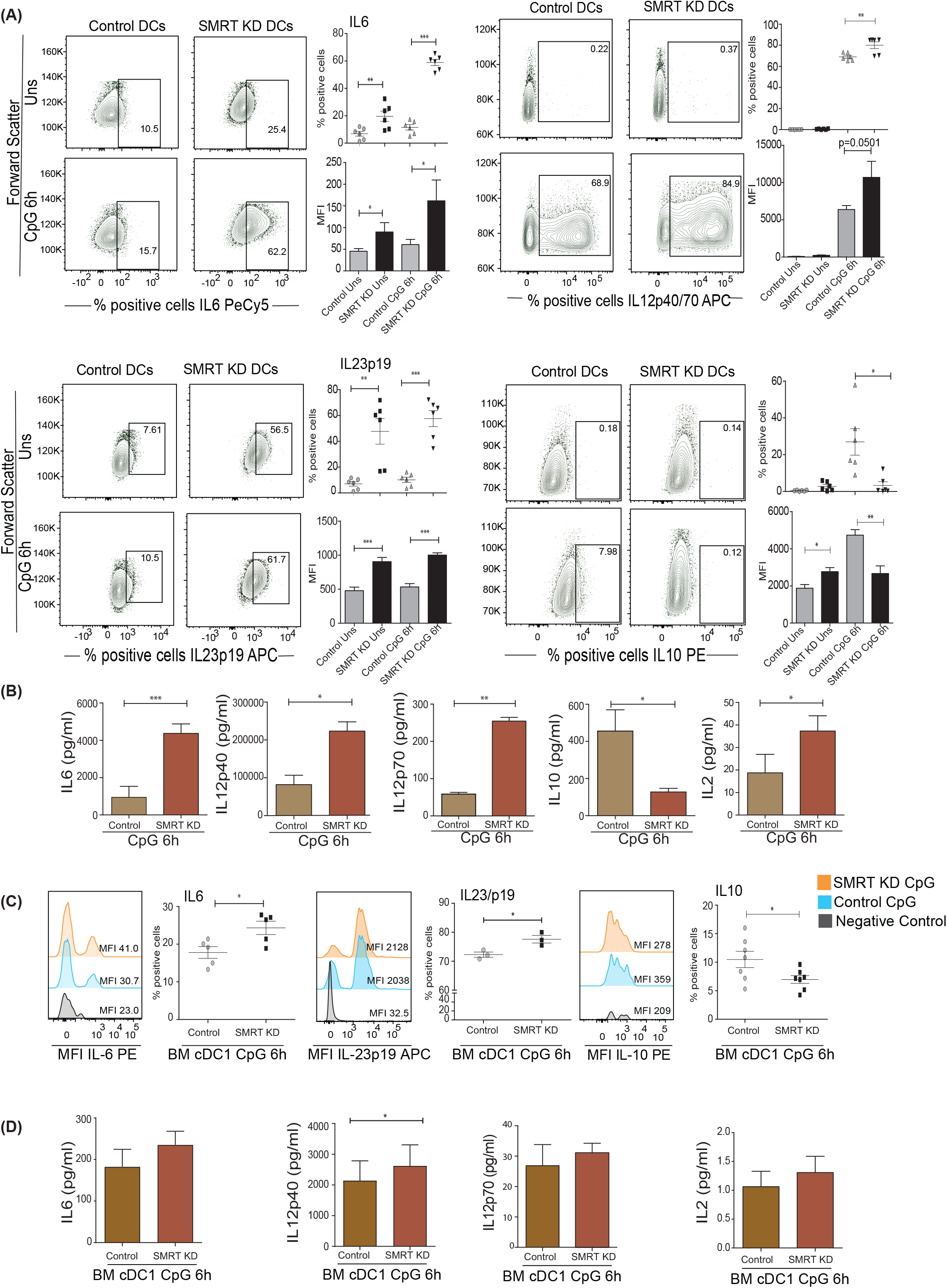
Activated SMRT KD cDC1 showed enhanced inflammatory cytokine expression. **A.** Flow cytometry analysis depicting the intracellular cytokine expression of IL-6, IL-12p40, IL-23p19, and IL-10 cytokine expression in control and SMRT KD cDC1 before and after 6h CpG challenge. Corresponding contour plot, dot plot and bar-plots show the percent positive cell population and MFI shifts respectively. (n=6) **B.** Bioplex cytokine assay showing the estimation of secreted cytokines IL-6, IL-12p40, IL-12p70, IL-10, and IL-2 in the culture supernatants of control and SMRT KD cDC1 before and after 6h CpG activation. (n=3-5) **C.** Flow cytometry analysis depicting the MFI histogram and scatter dot plots depicting the flow cytometry analysis for intra-cellular expression of pro-inflammatory cytokines IL-6, IL-23p19 and anti-inflammatory cytokine IL-10 in 6h CpG stimulated control and SMRT KD bone-marrow derived cDC1 (BMcDC1). (n=3-7) **D.** Bioplex cytokine assay showing the quantification of IL-6, IL-12p40, IL-12p70, and IL-2 cytokine secreted in the culture supernatant of 6h CpG stimulated control and SMRT KD BMcDC1s. (n=4-5) *p ≤ 0.05, **p ≤ 0.01 and ***p ≤0.001. p-value has been calculated using two tailed paired student’s t-test. Data shown in figure is combined from 3 independent experiments [A], 3 independent replicates [B], 2-4 independent replicates [C], and from 2 independent experiments [D]. Error bars represent SEM.

### Co-culture of OT-II CD4^+^ Th cells with SMRT depleted cDC1 enhanced Th1 and Th17 differentiation

To understand the functional impact of perturbed SMRT depleted DC responses on CD4^+^ Th cell differentiation, we performed a co-culture experiment of CD4^+^ Th cells isolated from OT-II transgenic mice with SMRT KD and control cDC1 mutuDC line. DCs were pulsed with OT-II peptide overnight, followed by activating with CpG or pIC for 2h before addition of OT-II T-cells. The isolated CD4^+^ OT-II were labelled with efluor-670 proliferation dye and co-cultured with OVA pulsed and CpG or pIC activated DCs for 72h. We first looked into the proliferation of co-cultured T-cells and found that in SMRT KD conditions there was enhanced T-cell proliferation compared to controls (**Figure 3A-3B and Supplementary Figure 5A**). Proliferation index of OT-II T cells co-cultured with SMRT KD and control DCs however did not show significant changes. Then we profiled the T-cell subtype patterns to look for its impact on differentiation. Further, it is known that IL-6 leads to suppression of forkhead box protein P3 (FOXP3) TF expression thus repressing Treg differentiation [27]. IL-6 along with IL-23 contributes to the development of Th17 cells by inducing RORγt expression [28]. Also, IL-12p70 is known to upregulate T-bet generating Th1 subtype. In our experiment, we found a significantly increased percentage of CD44^+^T-bet^+^IFN-γ^+^ as well as CD44^+^RORγt^+^IL-17^+^ cells supporting enhanced frequency of Th1 and Th17 subtype in activated SMRT KD cDC1 condition as compared to control DCs (**Figure 3C-3F**). A similar increase in CD44^+^T-bet^+^IFN-γ^+^ cells was observed in pIC stimulated condition as well (**Supplementary Figure 5B**). To further confirm the increased Th17 polarization, we checked IL-17 cytokine levels in the culture supernatants and found it to be significantly increased (**Figure 3G**). No significant change was observed in IL-13 secretion (**Supplementary Figure 5C**). There was no significant change observed for IL-10, Foxp3^+^ Tregs or GATA3^+^ Th2 cells (**Supplementary Figure 5D**). The gating strategy used for the analysis is depicted (**Supplementary Figure 5E**).

**Figure 3.**
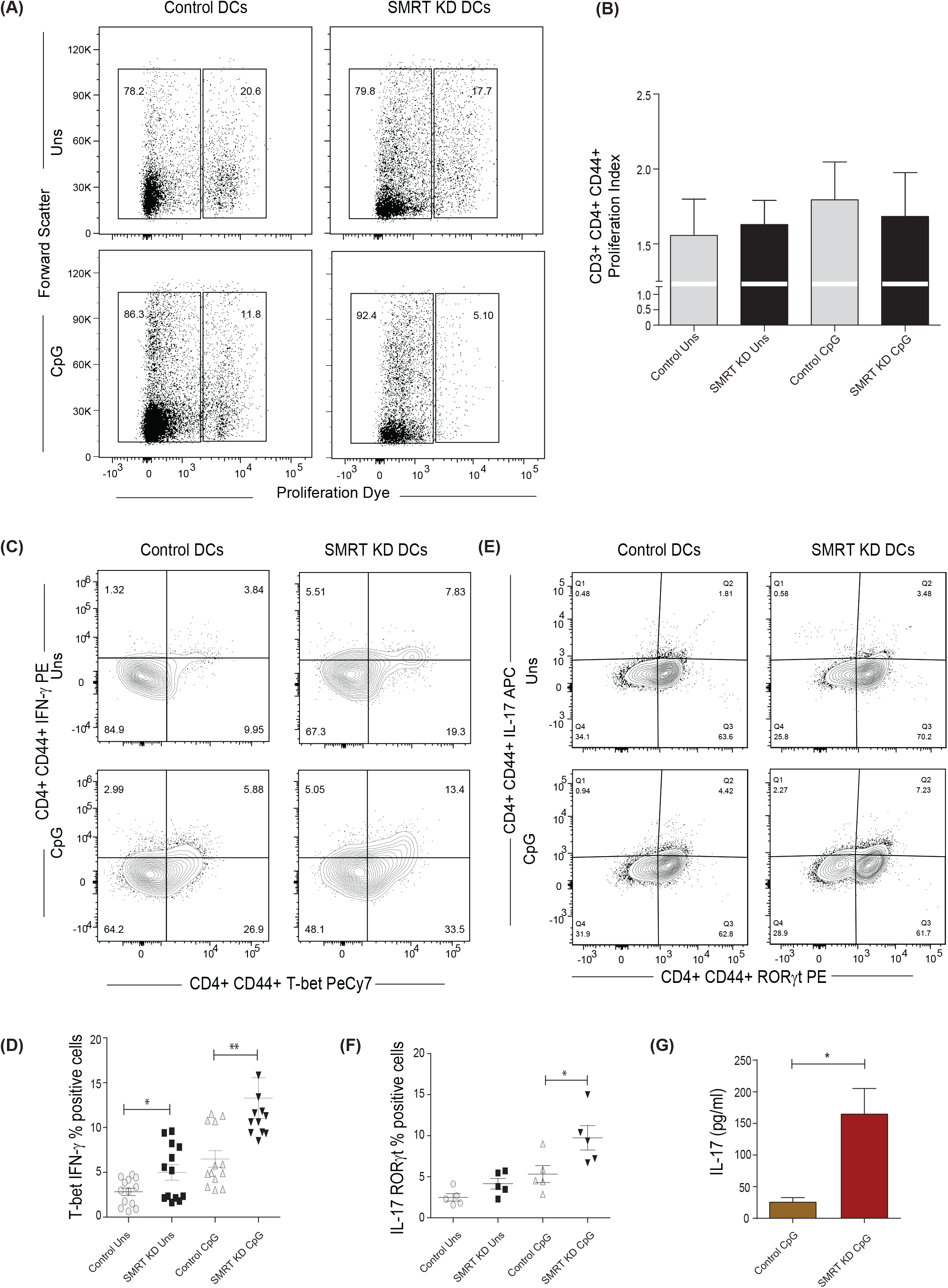
SMRT KD cDC1s enhanced Th1 and Th17 cell polarization *ex vivo*. **A.** Flow cytometry analysis depicting scatter plot to show the proliferating populations of OT-II Th-cells co-cultured with control and SMRT KD cDC1s pulsed with OVA 323-339 peptide overnight followed by CpG challenge. **B.** Flow cytometry analysis showing proliferation index of OT-II Th-cells co-cultured with control and SMRT KD cDC1s pulsed with OVA 323-339 peptide overnight followed by CpG challenge. (n=6) **C.** Flow cytometry analysis showing contour plots representing co-cultured OT-II T-cells showing signature transcription factor and cytokine for Th1, T-bet and IFN-γ response in unstimulated and CpG stimulated condition. **D.** Flow cytometry analysis showing scatter dot plots representing co-cultured OT-II T-cells showing T-bet and IFN-γ double positive cells in unstimulated and CpG stimulated condition. (n=13) **E.** Flow cytometry analysis showing contour plots showing RORγt and IL17A as signature transcription factor and cytokines for Th17 subtype generated from co-cultured OT-II T-cells in unstimulated and CpG stimulated condition. **F.** Flow cytometry analysis showing scatter dot plots showing RORγt and IL17A double positive cells for Th17 subtype generated from co-cultured OT-II T-cells in unstimulated and CpG stimulated condition. (n=5) **G.** Bioplex cytokine assay showing quantification of secretory cytokine IL-17 cytokine from supernatant of CD4^+^ T-cells co-cultured with CpG stimulated control and SMRT KD DCs. (n=5) *p ≤ 0.05, **p ≤ 0.01 and ***p ≤0.001. p-value has been calculated using two tailed unpaired student’s t-test. Data shown in figure is combined from 3 independent experiments [A-B], 3-5 independent replicates [D & F], and 3 independent experiments [G]. Error bars represent SEM.

### *Ncor2* expression is decreased in PBMCs of Rheumatoid Arthritis (RA) patients

The autoimmune diseases like Rheumatoid Arthritis (RA) have been widely classified as Th1 and Th17 disease and IL-10 expression is also found to be drastically reduced [29]. It had been established earlier that Th1 responses were involved in autoimmune disease. However, work on IFN-γ and IL-12^-/-^ mice showed that these mice have a high probability of developing collagen-induced arthritis. With this came the advent of Th17 cells and their role in autoimmunity [30]. Therefore, to identify if in such diseases *Ncor2* expression is decreased and if it correlates with inflammatory phenotype, we performed RT-qPCR of *Ncor2* from peripheral blood mononuclear cells (PBMCs) of 14 healthy donors and 11 RA patients. Our results demonstrated significantly lower *SMRT* expression in RA patients compared to their healthy counterparts (**Supplementary Figure 5F**). RA patients with disease activity score (DAS) above 4 have been considered in the study. This result suggested that *Ncor2* decrease could be associated with increased inflammatory phenotype in RA disease. All subjects provided informed consent and the study was approved by the Institutional Ethics Committee (KIIT/KIMS/IEC/39/2019).

### Co-culture of SMRT KD cDC1 with OT-I CD8^+^ T-cells increases T-cell cytotoxicity

As we observed significantly increased MHC-I expression on SMRT depleted DCs, we were interested to identify if these DCs have the potential to increase cytotoxic activity of CD8^+^ T-cells. For the same, CD8^+^ T-cells isolated from the spleen of OT-I transgenic mice were co-cultured with control and SMRT KD cDC1s. DCs were first pulsed with SIINFEKL (OVA peptide 257-264) with or without CpG or pIC for 2h followed by coculture with purified efluor-670 labelled CD8^+^ T-cells for 72h to assess proliferation and its impact on IFN-γ, perforin and granzyme-B (GrB) expression. No significant change was observed in the proliferation of co-cultured OT-I CD8+ T-cells with SMRT KD condition as compared to control DCs (**Figure 4A and 4B**). Activation profiling of cocultured CD8+ T-cells showed an increased frequency of IFN-γ, GrB, and Perforin expressing CD8+CD44+ T-cells in CpG activated SMRT depletion condition as compared to controls **(Figure 4C).** Similar results were obtained for GrB and Perforin using pIC stimulated conditions. IFN-γ showed a non-significant but increasing trend (**Supplementary Figure 5G**). The gating strategy used for the analysis is depicted (**Supplementary Figure 5H**).

**Figure 4.**
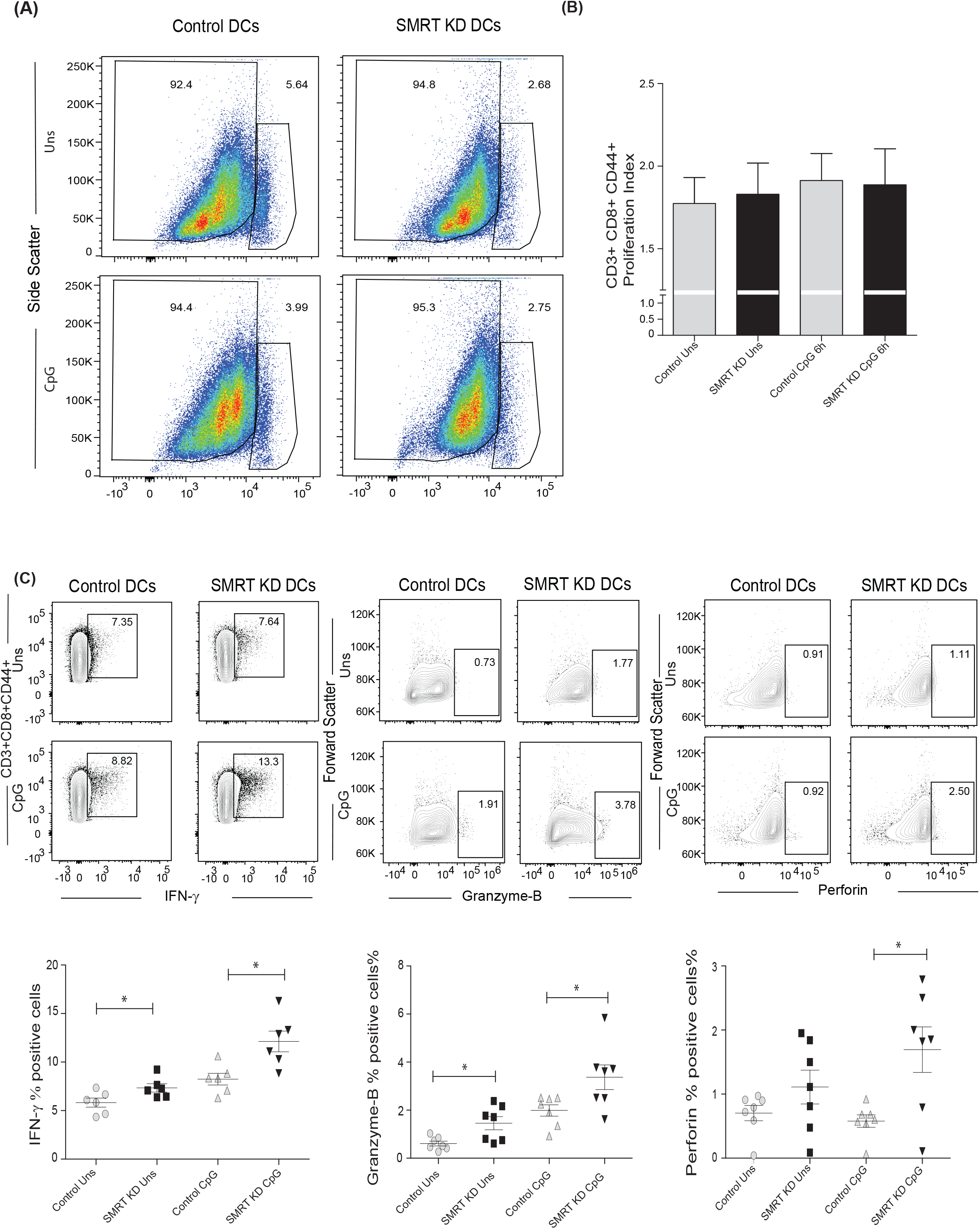
SMRT KD cDC1s increased perforin, granzyme, and IFN-γ production *ex vivo* in CD8 T lymphocytes. **A.** Flow cytometry analysis showing scatter plots of the proliferation of co-cultured OT-I T-cells co-cultured with control and SMRT KD cDC1s pulsed with OVA 257-264 peptide overnight followed by CpG challenge. **B.** Flow cytometry analysis showing the bar-plot demonstrating proliferation index of OT-I T-cells co-cultured with control and SMRT KD cDC1s pulsed with OVA peptide overnight followed by CpG challenge. (n=7) **C.** Flow cytometry analysis showing the contour plot and scatter dot plots representing co-cultured OT-I T-cells showing cytolytic IFN-γ, GrB, and Perforin in control and SMRT KD DCs in unstimulated and CpG stimulated condition. (n=6-7) *p ≤ 0.05, **p ≤ 0.01 and ***p ≤0.001. p-value has been calculated using two tailed unpaired student’s t-test. Data shown in figure is combined from 2 independent experiments [A & B], 2 independent replicates [C]. Error bars represent SEM.

### Adoptive transfer of OVA-Delayed Type Hypersensitivity (OVA-DTH) animals with CpG pulsed SMRT depleted cDC1 enhanced foot pad inflammation

OVA induced Delayed Type Hypersensitivity (OVA-DTH) has been employed to manifest antigen specific T-cell response which is dependent on efficiency of antigen presentation by cells in host where T-cell experienced antigen initially in a sensitization phase and recall response in their challenge phase later on [31]. We asked whether SMRT deficient DCs can elicit strong inflammatory antigen specific T cells response *in-vivo* as compared to its counterpart control cDC1 cells. To address our hypothesis, we developed the OVA-DTH mice model where we sensitize the mice with adjuvant emulsified OVA by subcutaneous (s.c) injection at day 0. Control and SMRT KD cDC1 pulsed with CpG and OVA were adoptively transferred at day 14. After one week, mice received OVA challenge locally in the left footpad to induce hypersensitivity mediated inflammation on day 20. Footpad inflammation was measured every 12h till 72h after the OVA challenge. After 72h, the animals were dissected to assess the impact of DC treatment on T-cell immune-modulation (**Figure 5A**). We found that treatment of SMRT depleted and activated cDC1 DCs in OVA sensitized animals showed profound and consistent increase in footpad inflammation till 72h (**Figure 5B and Supplementary Figure 6A**). Next, we examined T-cell subtypes in the popliteal and inguinal lymph nodes. Mice injected with SMRT KD DCs exhibited an increasing, although not significant, trend in Th1 subtype in lymph nodes as compared to control, and a significant increase compared to PBS treated group, marked by increased IFN-γ and T-bet positive T-cell population (**Figure 5C**). At the same time, we found a significantly higher Th17 T-cell population as evident from increased IL-17 and RORγt positive cells in mice injected with SMRT KD cDC1 (**Figure 5D**). It has been reported that Th1 response induces tolerance by inhibiting Th17 response which was consistent in control cDC1 treated mice [31]. This phenomenon was abolished in mice treated with SMRT KD cDC1 cells. Next, we asked the specificity of this observed T cell response by assaying the level of IgG subtypes in the sera of these mice collected at 0h and 72h from treated and control animals. We found that both mice groups treated with OVA pulsed control and SMRT cDC1 showed induced OVA specific IgG subtypes as compared to mice treated with PBS. We did not find any difference between OVA pulsed control and SMRT KD cDC1 treated mice group (**Supplementary Figure 6B**). These results showed that decreased SMRT level in DCs generated a very strong inflammatory T cells response in the host.

**Figure 5.**
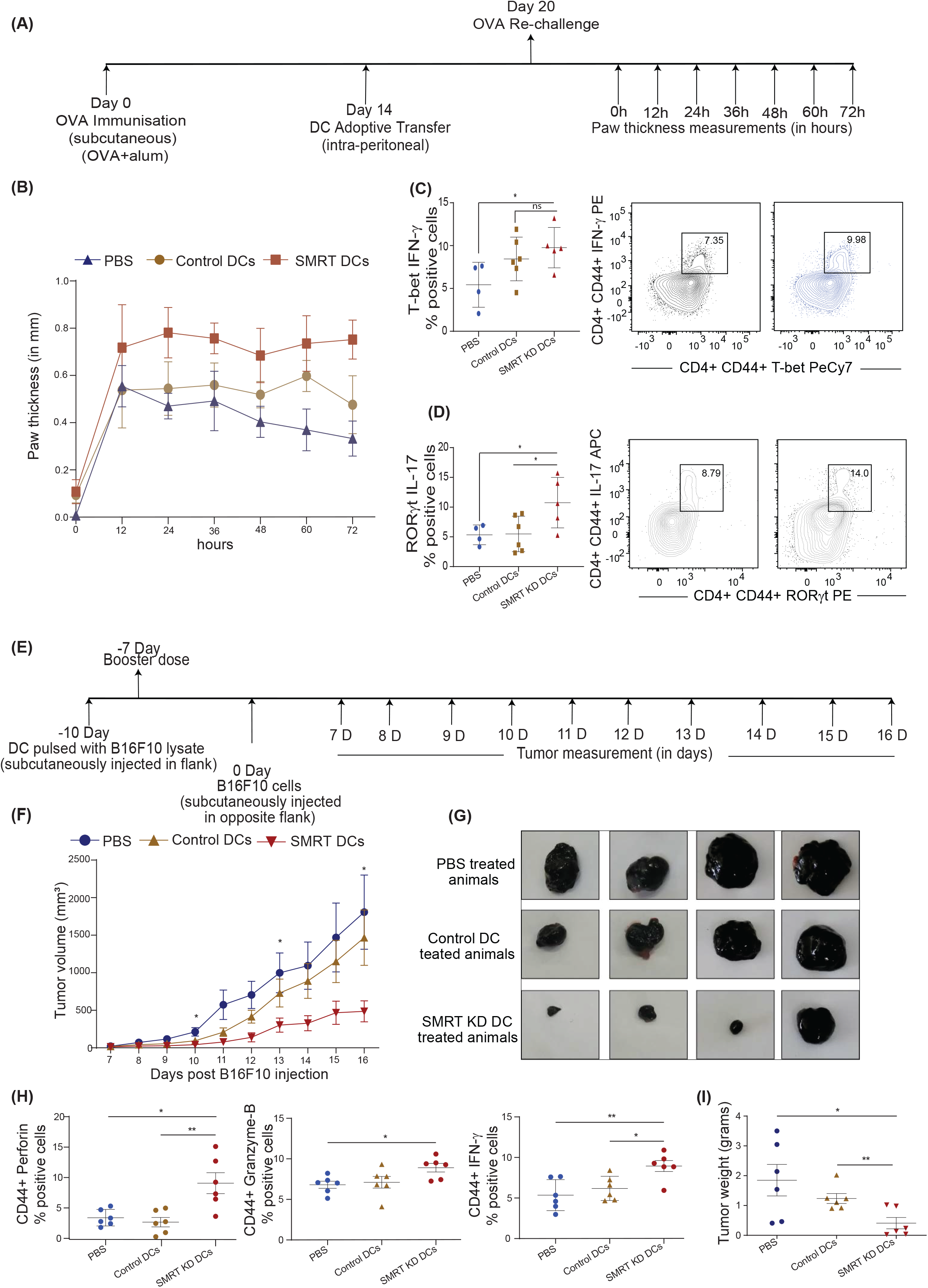
Induction of DTH response and tumor regression in C57BL/6 after adoptive transfer of Control and SMRT KD DCs. **A.** Experimental design depicting DTH model to understand the effect of SMRT KD in comparison to control cells through ova immunisation, sensitisation and rechallenge followed by measurement of paw thickness. **B.** Line plot with standard error mean showing footpad swelling till 72h post antigen rechallenge. Footpad swelling was calculated by subtracting the paw thickness (mm) in the right footpad (pbs injection) from left footpad (ova injection). (n=4-6) **C.** Flow cytometry analysis showing scatter dot and contour plot representing Th1 subtype marked by non-significant but enhanced IFN-γ and T-bet in SMRT KD DCs isolated from popliteal lymph nodes 72 h post antigen/ova rechallenge. (n=4-6) **D.** Flow cytometry analysis depicting scatter dot and contour plots representing Th17 subtype marked by enhanced IL-17 and RORγt in SMRT KD DCs isolated from popliteal lymph nodes 72 h post antigen/ova rechallenge. (n=4-6) **E.** Experimental design of melanoma model to understand the effect of SMRT KD DCs on tumor regression. Mice were first given tumor vaccination 10 and 7 days prior to tumor injection using DCs pulsed with B16F10 lysates. **F.** Line plot with standard error mean showing tumor volume that was taken every day starting from 7th day after B16F10 injection till 16 days post tumor rechallenge in PBS, control and SMRT KD DCs injected mice. Tumor volume was calculated as tumor volume = (tumor length x tumor width^2^)/2. (n=4-6) **G.** Images showing tumor size isolated from mice at 16 days post B16F10 rechallenge in PBS, control and SMRT KD DCs injected mice. (n=4) **H.** Flow cytometry analysis showing dot plots representing cytotoxic cytokine such as perforin, GrB, IFN-γ from single cell suspension of tumours that were isolated 16 days post B16F10 rechallenge in PBS, control and SMRT KD DCs injected mice. (n=6) **I.** Flow cytometry analysis showing dot plots showing tumor weight isolated 16 days post B16F10 rechallenge in PBS, control and SMRT KD DCs injected mice. (n=6) *p ≤ 0.05, **p ≤ 0.01 and ***p ≤0.001. p-value has been calculated using two tailed unpaired student’s t-test. Data shown in figure is combined from 2 independent experiments [A-I]. Error bars represent SEM.

### Adoptive transfer using CpG pulsed SMRT KD cDC1 regresses murine B16F10 melanoma tumor burden

It has been reported widely that DC vaccine therapy induces tumor associated antigen (TAA)-specific oncolytic CD8^+^ T-cell activity [32]. Therefore, to assess the physiological impact of enhanced cytotoxic CD8^+^ T-cell induced by SMRT depleted DCs, we developed a B16F10 melanoma model in C57BL/6 mice. The major objective was to check if SMRT KD cDC1 treatment could enhance and attract oncolytic CD8^+^ T-cell population in the developing tumor. First, we vaccinated the animals with B16F10 melanoma tumor antigen primed SMRT KD or control cDC1 to induce immunity against B16F10 tumors. We hypothesized that animals that received vaccination using inflammatory SMRT KD cDC1 loaded with B16 antigens could resist the increasing tumor burden compared to their control littermates. The vaccination with tumor antigen loaded SMRT KD or control cDC1 were done 10 days prior to subcutaneous injection of B16F10 cells for tumor development. To prime or load the control and SMRT KD cDC1 with B16F10 melanoma TAA, we pulsed control and SMRT KD DCs with B16F10 tumor lysate at 1:1 ratio prepared from equal number of B16F10 melanoma cells. Pulsing with tumor lysate was performed overnight for TAA presentation on DCs. Next day the TAA loaded DCs were washed and pulsed with CpG for 2h and 0.5 x 10^6^ cells were injected per mice in the shaved left flank of the mice. Mice were given vaccination in the form of TAA loaded DCs 10 days prior and a booster dose was given 7 days prior to tumor injection. Tumor development was induced in these mice after 7 days by injecting 0.1×10^6^ B16F10 cells subcutaneously in mice on the other flank (**Figure 5E**). Tumor volumes were measured every alternate day starting from 7 days till 16 days post tumor initiation and we found that in the SMRT cDC1 treated group the tumor burden was significantly reduced as compared to control groups **(Figure 5F, 5G and Supplementary Figure 6C).** Further to assess the CD8^+^ T-cell cytotoxicity, we sacrificed the mice at 16 days post tumor initiation, dissected the tumors and made single cell suspension by collagenase treatment. The cells were then stimulated using PMA/Ionomycin/BFA for 5h, and CD3^+^CD8^+^ T-cells were analyzed. We found that frequency of Perforin, GrB and IFN-γ percentage positive cells were significantly increased in SMRT KD cDC1 vaccinated animals as compared to other treatment groups **(Figure 5H and Supplementary Figure 6D, Supplementary Figure 6E)**. The tumors isolated from all the three groups of the animals showed a significantly decreased tumor weight in SMRT KD cDC1 vaccinated condition **(Figure 5I).**

### Comparative genomic and transcriptomic analysis of NCoR1 and SMRT in cDC1 showed differential control of STAT3 signaling mediated IL-10 regulation

Recently, we reported that NCoR1 directly binds and strongly represses tolerogenic genes such as *Il10, Il27, Cd83* and *Socs3* in cDC1. NCoR1 depletion drastically increased the expression of these genes thereby leading to Treg generation *ex vivo* and *in vivo* [25]. Contrary to this, here we found that NCoR1 paralog SMRT depleted cDC1 demonstrated drastically increased inflammatory response *in vitro, ex vivo* and *in vivo*. Therefore, we performed comparative genome-wide binding of NCoR1 and SMRT and transcriptomic analysis of NCoR1 and SMRT KD DCs to understand the differential gene regulation by NCoR1 and SMRT. Differential gene expression analysis of SMRT KD versus control cDC1 at 6h CpG stimulation showed 1273 and 934 genes up- and down-regulated respectively (Log2 Fold change >1 and adjusted p-value ≤ 0.01) (**Figure 6A, Supplementary File 1**). Large number of direct target genes identified using ChIP-seq analysis of SMRT that are upregulated compared to downregulated support its role as a co-repressor (**Figure 6B, Supplementary File 2**). Further, Ingenuity pathway analysis (IPA) of direct target differentially up regulated genes in SMRT KD condition depicted up-regulation of inflammatory response pathways such as IL-12 signaling (**Figure 6C, Supplementary File 3**). However, in contrast to NCoR1, SMRT KD showed downregulation of IL-10 signaling pathway at 6h CpG stimulation (**Figure 6D, Supplementary File 3**). Next, principal component analysis of NCoR1 and SMRT KD cDC1 RNA-seq clearly showed SMRT KD unstimulated and 6h CpG stimulated condition cluster separately from respective NCoR1 samples (**Supplementary Figure 7A**). Interestingly, in unstimulated condition the number of differentially regulated genes (up-regulated (n=1060) and down-regulated (n=805)) are much higher in SMRT KD condition as compared to NCoR1 KD DCs (**Supplementary Figure 7B-C, Supplementary File 1**). The PCA analysis and difference in the number of differentially expressed genes (DEGs) between SMRT and NCoR1 KD indicated a differential control of gene regulation by SMRT and NCoR1 in cDC1. To identify gene sets showing different pattern of expression after NCoR1 and SMRT KD, we performed unsupervised K-means clustering of DEGs based on log2 fold change and identified six clusters (**Figure 6E and Supplementary Figure 7D, Supplementary File 4**). The genes in cluster-1 showed increased fold change in both NCoR1 and SMRT KD and depicted enriched pathways such as “Interferon signaling”, “Activation of IRF by cytosolic pattern recognition receptors”, “IL-12 signaling and production in macrophages”. On the contrary, cluster-2 genes showed differential regulation between SMRT and NCoR1 KD 6h CpG condition and pathways enriched were “Th2 pathway”, “STAT3 pathway”, “IL-17-A signaling” and “IL-10 Signaling”. (**Figure 6F, Supplementary File 5**). Other clusters (3-6) genes also showing similar or differential effect upon NCoR1 and SMRT KD were enriched for cytokine signaling pathways terms such as “IL-8”, “IL-7” and “IL-15” signaling (**Supplementary Figure 7D-E, Supplementary File 2**). These observations clearly suggested that inflammatory immunogenic response genes such as *Il12b* and *Il6* are repressed by both NCoR1 and SMRT, however, the regulatory genes like *Il10, Socs3* are strongly repressed by NCoR1 only (**Figure 6G**). Apart from the above listed gene, we also found other reported positive regulator of pro-inflammatory genes such as *Zbtb20* is upregulated and positive regulator of tolerogenic program such as *Wnt11, Clec4a2* is downregulated in SMRT KD cDC1s (**Supplementary File 1**) [33–36]. As IL-10 is a physiologically important cytokine and well reported to perturb the inflammatory response towards immune-tolerance, understanding mechanistic control of differential IL-10 regulation by SMRT and NCoR1 was interesting [37]. We compared the cistrome of SMRT and NCoR1 before and after 6h CpG activation (~13,000 peaks) (**Supplementary Figure 8A**). In SMRT, 40% (≈5000) of the total bound sites are distributed at the promoter-proximal (± 1kb to transcriptional start site (TSS)) regions whereas only 18% of NCoR1 peaks (≈2000 peaks) were found at promoter-proximal regions (**Supplementary Figure 8B**). Besides, differential binding of NCoR1 and SMRT peaks identified NCoR1 dominant (12,473), common NCoR1-SMRT (5949) and SMRT dominant (2707) genomic regions (**Figure 6H, 6I and Supplementary Figure 8C**). SMRT dominant regions showed predominance in promoter-proximal regions compared to NCoR1 dominant and common NCoR1-SMRT regions distributed mostly in intronic or distal intergenic regions (**Figure 6H, and Supplementary Figure 8C**). This further strongly suggested that SMRT mostly regulate the genes through promoter-proximal binding, whereas NCoR1 and common NCoR1-SMRT bound genes are regulated through far-distal regulatory elements. Further, KEGG pathway analysis of genes associated with genomic regions showing CpG dependent increase in NCoR1 or common NCoR1-SMRT binding is significantly enriched for “Th1 and Th2 pathway”, “Th-17 signaling pathway”, “NFkB signaling pathway”, and “JAK-STAT signaling pathway” (**Figure 6J and Supplementary Figure 8D, Supplementary File 6**). Co-occupancy of NCoR1 as well as SMRT on both inflammatory (*Il12b*) and tolerogenic genes (*Il10*) at several proximal or distal regions to TSS after CpG activation shows involvement of both the repressor in regulating immune response genes (**Figure 6K**). To identify the putative transcription factors (TFs) recruited before and after 6h CpG activation at these co-repressors bound genomic regions, we performed de novo motif enrichment analysis and found that PU.1, RUNX2 and Jun-Fos/AP1 TFs motifs were enriched at almost all NCoR1 and SMRT bound genomic regions showing enrichment in unstimulated or 6h CpG activation condition. Interestingly, NFkB motifs were found to be enriched at CpG dominant NCoR1, dominant SMRT and common NCoR1-SMRT binding. IRF8 and IRF4 motifs were enriched at dominant NCoR1 and common NCoR1-SMRT regions in both unstimulated and 6h CpG activation conditions respectively (**Figure 6L and Supplementary Figure 8E**). The above enriched motifs corroborate with our previous finding on NCoR1 bound regions [25]. Apart from the above motifs we found an enrichment of STAT3 motif at CpG dominant common NCoR1-SMRT and CpG dominant SMRT genomic regions. Since member of NFkB family and Stat3 transcription factor plays an important role in regulation of inflammatory (*Il12b, Il6*) and tolerogenic (*Il10, Socs3*) gene expression, to further understand downregulation of tolerogenic genes, we checked expression of other genes associated with Jak-Stat signaling pathway. We found a clear difference in regulation of several other Jak-Stat signaling which is downregulated in SMRT KD DCs but not in NCoR1 KD DCs (**Figure 7A, 7B**). Overall, our comparative genomic and transcriptomic analysis suggest that SMRT KD DCs show dysregulation of STAT3-IL-10 axis in contrast to NCoR1 KD DCs.

**Figure 6.**
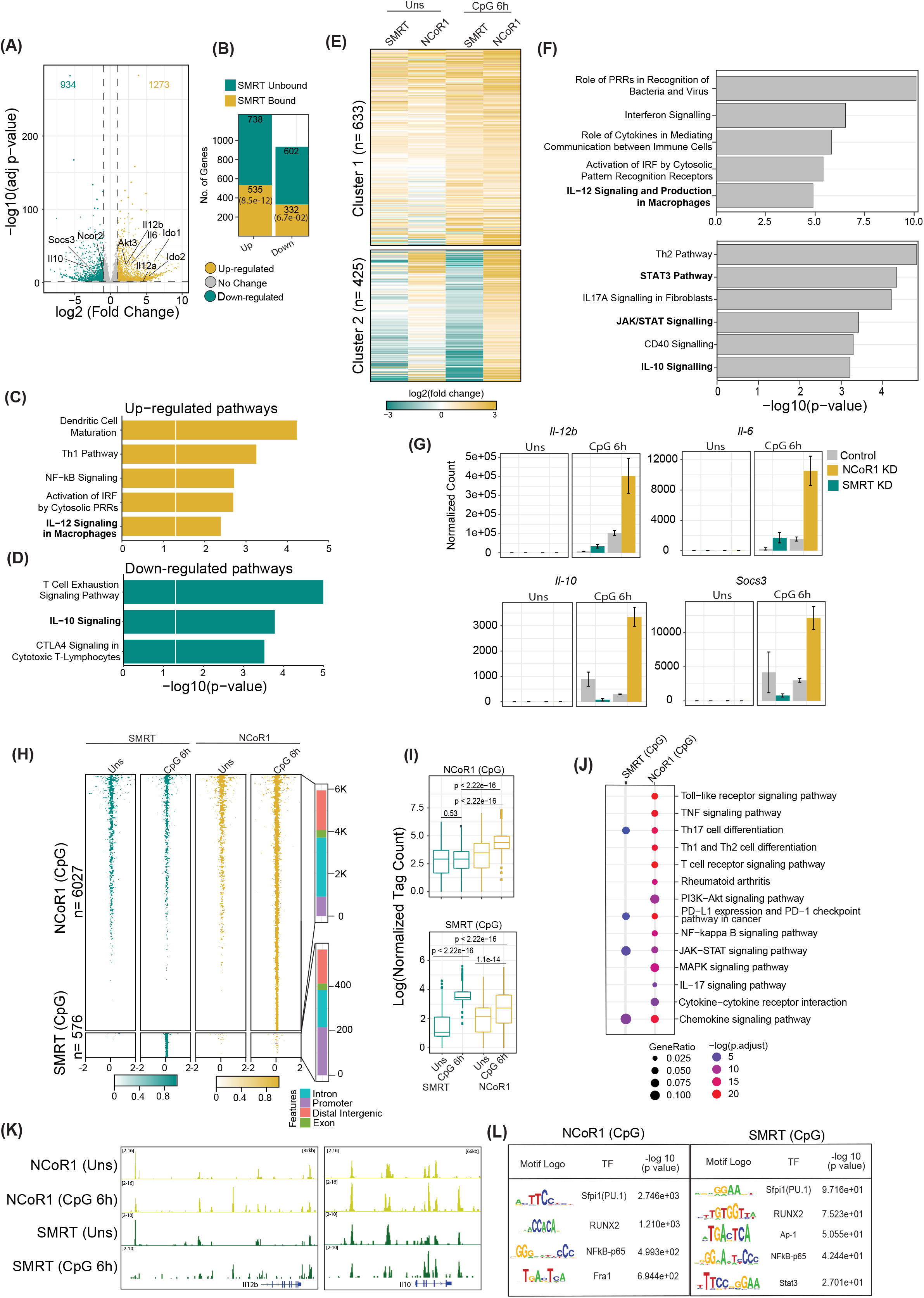
Integrative genomics analysis (RNA-seq and ChIP-seq) identified the differential role of SMRT and NCoR1 in regulation of immune response in cDC1. **A.** Volcano plot showing the differentially expressed genes (DEGs) in SMRT KD cDC1 DCs as compared to control cells after 6h CpG stimulation. There were 1273 and 934 genes upregulated and downregulated respectively upon SMRT depletion. (n=5) **B.** Bar plot showing the total number of SMRT bound and unbound DEGs in unstimulated and 6h CpG stimulated SMRT KD cDC1 DCs. p-value showing significance of overlap between SMRT bound genes and DEGs. **C.** Bar plot depicting enriched canonical pathways for the list of SMRT bound upregulated DEGs in 6h CpG activated SMRT KD cDC1 DCs as compared to control cells using Ingenuity pathway analysis (IPA). **D.** Bar plot depicting enriched canonical pathways for the list of significantly SMRT bound down-regulated DEGs in 6h CpG activated SMRT KD cDC1 DCs as compared to control cells using Ingenuity pathway analysis (IPA). **E.** Heatmap showing two clusters obtained from K-means clustering of log2 fold change of DEGs upon SMRT and NCoR1 KD compared to control cells in unstimulated and 6h CpG stimulation condition. **F.** Bar plot showing enriched canonical pathways from IPA for respective clusters shown in Fig 6E. **G.** Bar plot with standard deviation showing normalized count from DESeq2 of selected genes from enriched pathway for cluster-1 (*Il12b* and *Il6*) and cluster2 (*Il10* and *Socs3*). **H.** Tornado plot showing ChIP-seq signal (±2kb to peak center) of differential NCoR1 and SMRT binding sites (NCoR1 CpG and SMRT CpG) in unstimulated and 6h CpG stimulation (1^st^ panel). Bar plot showing the distribution of differential genomic regions based on distance relative to TSS (2^nd^ panel). **I.** Box plot showing normalized tag count in differential NCoR1 CpG and SMRT CpG cluster. Boxes encompass the 25^th^ to 75^th^ percentile of normalized tag count. Whiskers extend to the 10^th^ and 90^th^ percentiles. Mean difference significance was calculated using the Wilcoxon test. **J.** Dot plot showing the significantly enriched KEGG terms for genes associated with NCoR1 and SMRT CpG binding clusters shown in Fig 6I. KEGG term enrichment analysis was performed using cluterProfiler R package. **K.** IGV snapshot showing NCoR1 and SMRT binding on IL12b and Il10 gene loci. **L.** Table showing transcription factor motifs that were significantly enriched (P-value < 1e-10) in differential NCoR1 CpG and SMRT CpG clusters.

**Figure 7.**
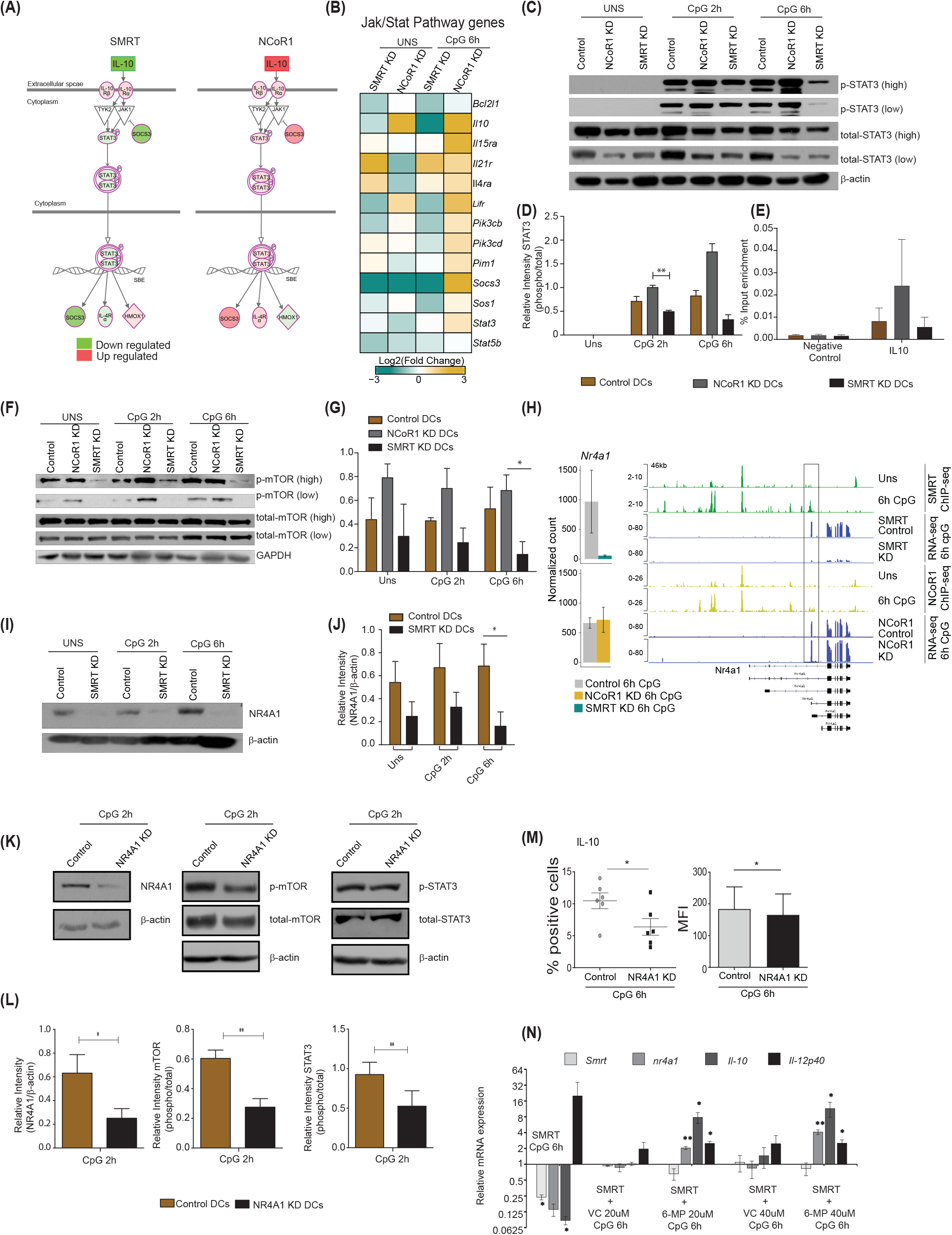
Nurr-77, mTOR, Stat3 signaling regulates IL-10 expression in SMRT KD cDCs. **A.** Illustration of Jak/Stat Signaling pathway from Ingenuity pathway analysis with mapped expression of genes. (Red and green indicate high and low expression respectively). **B.** RNA-seq analysis showing heatmap showing Log2 (fold change) of all the genes in KEGG Jak-Stat signaling term shown in Fig 6L. **C.** Western blot depicting phospho-STAT3, total STAT3, and β-actin protein levels in unstimulated and 2h and 6h CpG stimulated control, NCoR1 KD, and SMRT KD cDC1s. **D.** Bar plot with standard error mean from densitometric analysis depicting normalized intensity of phosphorylated STAT3 bands in control and KD cDC1s. Housekeeping gene β-actin was used as loading control. (n=3). **E.** Bar plot with standard error mean showing % input enrichment from ChIP-qPCR of phospho-STAT3 on IL-10 enhancer region in 2h CpG stimulated control, NCoR1 KD, and SMRT KD cDC1s. (n=3) **F.** Western blot depicting the levels of phosphorylated mTOR, total mTOR, and GAPDH in unstimulated, 2h, and 6h CpG stimulated control, NCoR1 KD, and SMRT KD cDC1s. Housekeeping gene GAPDH was used as loading control. **G.** Bar plot with standard error mean from densitometric analysis depicting normalized intensity of phosphorylated mTOR bands in control and KD cDC1s. Housekeeping gene GAPDH was used as loading control. (n=3) **H.** Bar plot with standard deviation showing normalized count of *Nr4a1* gene in NCoR1 and SMRT KD cDC1 with their respective matched controls in 6h CpG stimulation. IGV snapshot showing SMRT and NCoR1 binding at *Nr4a1* gene loci in control unstimulated and 6h CpG stimulated DCs along with RNA-seq in control, SMRT KD and NCoR1 KD DCs in 6h CpG stimulation condition. **I.** Western blot depicting the levels of NR4A1 and β-actin in unstimulated, 2h, and 6h CpG stimulated control and SMRT KD cDC1s. (n=3) **J.** Bar plot with standard error mean from densitometric analysis depicting the normalized intensity of NR4A1 bands in KD and control cells. Housekeeping gene β-actin was used as loading control. (n=3). **K.** Western blot depicting the levels of phospho-mTOR, total mTOR, phospho-STAT3, total STAT3, NR4A1, and β -actin in 2h CpG stimulated control and NR4A1 KD cDC1s. **L.** Bar plot with standard error mean from densitometric analysis depicting the normalized intensity of phosphorylated-mTOR, phosphorylated-STAT3, and NR4A1 in 2h CpG stimulated control and NR4A1 KD cDC1s. Housekeeping gene β-actin was used as loading control. (n=4) **M.** Percent positive cells and MFI depicting flow cytometry analysis of the anti-inflammatory cytokine IL-10 in 6h CpG stimulated control and NR4A1 KD DCs. (n=6) **N.** Relative transcript expression of *Ncor2, Nr4a1, Il10*, and *Il12b* transcript in 6h CpG-B stimulated SMRT KD, CpG along with vehicle treated SMRT KD and CpG along with 6-MP treated SMRT KD DCs as estimated by RT-qPCR. (n=3) *p ≤ 0.05, **p ≤ 0.01 and ***p ≤0.001. p-value has been calculated using two tailed paired student’s t-test. Data shown in figure is combined from 3-4 independent experiments [C-L], 4 independent experiments [M], and 3 independent replicates [N]. Error bars represent SEM.

### SMRT mediated down-regulation of NR4A1 inhibited mTOR-STAT3 signaling leading to IL-10 suppression

STAT3 TF plays a central role in Jak-Stat signaling and regulated expression of *Il10* and *Socs3*. We also found NFkB inhibitory genes such as *Nfkbia* and *Tnfaip3* were also downregulated after 6h CpG stimulation in SMRT KD cDC1. First to confirm the differential regulation of p-STAT3 in NCoR1 and SMRT, we did western blotting for p-STAT3 in NCoR1 KD, SMRT KD and control cDC1 at 0h, 2h, and 6h after CpG activation. We found that p-STAT3 is down-regulated in SMRT KD cDC1 compared to control cells whereas it was upregulated in NCoR1 depleted DCs (**Figure 7C-7D**). It is well reported that STAT3 binds to the *Il10* gene to regulate its expression [38]. We also checked STAT3 binding on *Il10* gene at 0h and 6h LPS stimulation in BMDCs (**Supplementary Figure 8F**) [39]. Therefore, we performed chromatin immunoprecipitation (ChIP) for p-STAT3 followed by RT-qPCR to infer the binding of p-STAT3 on *Il10* gene after 2h CpG challenge in control, NCoR1 KD, and SMRT KD cDC1. We observed reduction in p-STAT3 binding on *Il10* in SMRT KD DCs relative to control DCs whereas on the other side, the binding was found to be enhanced in NCoR1 KD cells as compared to control cells (**Figure 7E**). Moreover, to understand the upstream control of STAT3 signaling in these DCs we looked into the literature and identified that mTOR has been reported to control STAT3 activation and mTOR on the other hand is regulated by nuclear receptor NR4A1 also known as NURR-77 [40, 41]. We first checked the regulation of phospho-mTOR (p-mTOR) in NCoR1 and SMRT depleted cDC1 and found that p-mTOR is upregulated in NCoR1 KD cDC1 whereas it is drastically reduced after SMRT depletion (**Figure 7F, 7G**). In addition, we found differential regulation of *Nr4a1* (Nur-77) in SMRT and NCoR1 depleted DCs. SMRT KD leads to downregulation of *Nr4a1* while NCoR1 KD has no significant effect on expression of *Nr4a1* (**Figure 7H**). Further we checked direct binding of NCoR1 and SMRT on *Nr4a1* in ChIP-seq data using IGV browser and observed that SMRT but not NCoR1 binds at the TSS of the transcript that is expressed in DCs after 6h CpG activation (**Figure 7H**). We also confirmed this observation by assessing the NR4A1 protein expression in SMRT depleted and control cDC1 before and after 2h and 6h CpG activation and found that NR4A1 is significantly down-regulated in SMRT KD DCs as compared to control cells (**Figure 7I, 7J**). Further to confirm that NR4A1 indeed is controlling the STAT3 signaling through mTOR in SMRT depleted cells, we generated a stable NR4A1 KD and empty vector transduced (control) cDC1 using NR4A1 lentiviral shRNA to confirm its role in mTOR-Stat3-Il-10 signaling. In stable NR4A1 depleted mutu-cDC1 we first confirmed the depletion of NR4A1 by western blotting and found it to be significantly reduced **(Figure 7K, 7L).** Then we checked p-mTOR and phospho-STAT3 (p-STAT3) after 2h CpG activation which were found to be significantly reduced in NR4A1 depleted cDC1 compared to control cells **(Figure 7K, 7L)**. Moreover, in these NR4A1 depleted DCs we also confirmed significant reduction of IL-10 percent positive cells and corresponding MFI shifts upon 6h CpG treatment **(Figure 7M).** Moreover, we also tried to over-express NR4A1 ORF transiently in SMRT depleted cDC1 but we didn’t manage to perform this analysis as the cells were not in good condition after transduction of overexpression plasmids. Therefore, we used an alternative approach. It has been reported that 6-mercaptopurine (6-MP) induces the expression of NR4A1 in cells [20]. Therefore, we used 6-MP to enhance the expression of *Nr4a1* in SMRT depleted cDC1 to see if it can complement the *Il10* expression. We found that 6-MP treatment enhanced the expression of *Nr4a1* in SMRT depleted cDC1 leading to an increase in expression of *Il10* (**Figure 7N**). These results confirmed that SMRT mediated down-regulation of NR4A1 resulted in reduction of STAT3 activation and thereby decreased IL-10 levels and enhanced inflammatory phenotype of cDC1.

## Discussion

Dendritic cells make a strong connecting link between innate and adaptive immunity. Besides, a fine balance of DC signals is pertinent for development of an optimal immunogenic versus tolerogenic response to protect from autoimmune diseases and opportunistic infections. Though signaling pathways controlling one response versus another have been widely explored, how this balance is fine-tuned in DCs is interesting to understand for developing DC based therapies. We recently reported that NCoR1 directly binds and represses the transcription of regulatory genes like *Il10, Cd274, Cd83* and *Il27* and its loss of function in DCs enhanced Tregs development [25]. In contrast, SMRT depletion in cDC1 showed enhanced Th1 and Th17 frequency along with increased cytotoxic T-cell through increase in expression of pro-inflammatory genes (*Il12a, I12b, Il6, Il23a*) and decreased of tolerogenic genes such as *Il10* and *Socs3*. It has been demonstrated by Li. *et. al*. that macrophage specific depletion of NCoR1 in high-fat induced obese mice derepresses LXRs which generates anti-inflammatory response through increased expression of genes involved in fatty acid biosynthesis [42]. This is similar to the observations we have also made in cDC1s. The paradoxical effect of two paralog proteins, NCoR1 and SMRT, has been reported to be dependent on other interacting proteins in the co-repressor complex. As demonstrated by *Fan et. al*, the differential role of NCoR1 and SMRT is attributed to GPS2, a component of the corepressor complex which also contains HDAC3, NCoR1, SMRT, TBL1, and TBLR1 [43]. Macrophage specific deletion of GPS2 or SMRT leads to increased pro-inflammatory gene expression in macrophages while NCoR or HDAC3 depletion leads to enhanced anti-inflammatory effect [44, 45]. An extremely crucial observation in this study was that SMRT but not NCoR1 depletion hampered GSP2 recruitment to its target gene Ccl2, an inflammatory chemokine. Thus, GSP2 has been linked to SMRT, but not NCoR1, functionally which provides the inflammatory state in macrophages in GSP2 specific macrophage KO mice. These results support the fact that NCoR1 and SMRT are crucial determinants of inflammatory versus anti-inflammatory responses in immune cells. These findings depicted the role of these two co-regulators in fine-tuning inflammatory versus tolerogenic signals in cDC1 DCs. To understand this further, we performed comparative genomic analysis of NCoR1 and SMRT depleted DCs and found that they differentially regulate mTOR-STAT3 signaling pathway leading to tight regulation of IL-10. Overall, our analysis for the first time identified a fine switch that could be targeted to modulate inflammatory versus tolerogenic program in cDC1 DCs.

IL-10 is a potent anti-inflammatory cytokine that can limit host inflammatory response to pathogens thereby preventing host from damage. Dysregulated IL-10 is involved in enhanced immune-pathology and associated with development of auto-immune diseases [46]. The expression of IL-10 is dependent on STAT3 TF and it has been reported that STAT3 regulates *Il10* expression by binding to its regulatory region [13, 47]. At the same time, the positive feedback loop from IL-10 through STAT3 maintains its sustained expression. Immune-profiling analysis of NCoR1 and SMRT depleted cDC1 depicted significant upregulation of IL-10 in NCoR1, whereas it was drastically down-regulated in SMRT KD. When we looked at STAT3 regulation employing the integrative genomic analysis, it clearly showed significant down-regulation of STAT3 pathway in SMRT KD cDC1s. In contrast to this it was significantly upregulated in NCoR1 KD condition. Moreover, we found that there is increased binding of STAT3 on *Il10* in NCoR1 as compared to SMRT depleted cells. Overall, it is quite intriguing that two highly homologous co-regulators showed this differential regulation of an important pathway i.e., STAT3 and thereby IL-10 expression. Apart from STAT3 there are other transcription factors like Sp1/ Sp3, NF-kB, c-Maf, Smad4 that also exhibit a similar phenomenon in macrophages. However, studies showed that there is no increased binding of Sp1/Sp3 on *Il10* promoter in murine BMDCs [48]. We further explored to identify the mechanisms for differential regulation of STAT3 signaling. It has been reported that mTOR activates STAT3 [49] and we found that p-mTOR is up-regulated in activated NCoR1 KD cDC1 whereas it is drastically reduced in unstimulated as well as activated SMRT KD cDC1.

Furthermore, we observed that SMRT depleted DCs have sustained and increased expression of pro-inflammatory cytokines like IL-6, IL-12 and IL-23 thereby leading to enhanced development of Th1 and Th17 cells *ex vivo* and *in vivo* animal models. When we analyzed our global transcriptome data, we found that *Socs3* is significantly downregulated along with negative regulators of NFkB signaling such as *Nfkbia* and *Tnfaip3*. It has been widely reported that SOCS3 depletion enhances the Th1 and Th17 cytokine release in DCs [50]. The *Socs3* down-regulation was observed only in activated SMRT KD cDC1 after 6h activation, which suggested that the initial events of IL-10 and STAT3 decrease through down-regulation of mTOR activity are somehow leading to SOCS3 decrease. To our surprise, we found that SOCS3 and STAT3 both are down regulated in SMRT KD cDC1, as in several reports it has been documented that decreased SOCS3 favors enhanced STAT3 [51]. We hypothesize that it could be due to the dynamic time dependent regulation of STAT3 and SOCS3 in CpG activated SMRT KD cells.

Moreover, as we observed a drastic decrease of p-mTOR in unstimulated SMRT KD cDC1s, we looked into the regulators of mTOR. We found an interesting nuclear receptor *Nr4a1* to be significantly down-regulated in RNA-seq data of unstimulated SMRT depleted DCs and even after CpG activation. NR4A1 has been reported to positively regulate mTOR activity and thereby STAT3 phosphorylation and IL-10 expression [52]. It is also shown that deficiency of NR4A1, a member of the nuclear receptor superfamily, leads to enhanced production of IL-6, TNF-α, and IL-12 in both human and murine dendritic cells [20]. Similarly, peroxisome proliferator-activated receptor (PPAR-α) also exerts anti-inflammatory effects on monocytes and macrophages [53]. We showed that indeed NR4A1 expression was down-regulated supporting our finding in SMRT depleted DCs. It has been reported that ASC-2 and SMRT lead to transactivation and repression of Nr4A1 respectively [21]. We looked into the expression of ASC-2 (*Ncoa6*) and CaMKIV (*Camk4*) transcripts in SMRT KD cells but surprisingly we didn’t observe any significant change or downregulation. As we observed that SMRT KD downregulated a large number of genes even in unstimulated condition, it is plausible that an important Nr4A1 transactivation factor is also down-regulated leading to its decreased expression. On the other side, complementing *Nr4a1* in SMRT depleted cDC1 by treating with 6-MP rescued *Il10* expression. These indicated the contrasting inflammatory versus tolerogenic phenotype expressed by SMRT and NCoR1 depleted DCs.

In an *in vivo* physiological system, the maintenance of Th1, Th17, and Th2 balance depends on a number of factors including antigen presentation by MHC-II, costimulation and differential cytokine production [54]. To address whether the manipulation of DCs by SMRT KD towards an enhanced Th1 and Th17 type responses could deliver a long term functional and effector memory response, we developed DTH and B16F10 melanoma models. The SMRT depleted and control DCs were adoptively transferred and perturbation in DTH responses and B16F10 melanoma progression was observed. We found that SMRT depleted DCs have potential to enhance Th1 and Th17 type responses and thereby DTH enhancement in animals. At the same time, oncolytic CD8^+^ T-cell activity was enhanced leading to reduced tumor burden in animals. Furthermore, it has been widely reported that in autoimmune diseases such as RA and Multiple Sclerosis (MS) enhanced Th1 and Th17 cells result in inflammatory symptoms [55]. As we observed enhanced Th1 and Th17 response by SMRT depleted DCs we found significantly reduced SMRT expression in mononuclear cells of RA patients as compared to controls. Therefore, it is further interesting to explore if SMRT co-repressor has some potential association with autoimmune pathogenesis and can be used as a target for immunotherapy.

## Materials and Methods

### Mice

C57BL/6 wild type mice bred and maintained at ILS animal facility. OT-II and OT-I transgenic mice (gifted by Prof. Hans Acha-orbea, University of Lausanne) and C57BL6 Flt3 transgenic mice (gifted by Ton Rolink) were transported from SWISS. All the animal experiments were performed after getting due approval from the institutional animal ethics committee (ILS/IAEC-164-AH/AUG-19) and (ILS/IAEC-123-AH/AUG-18).

### Cell Lines

The CD8α+ mutuDC cell line used in this study has been gifted by Prof. Hans Acha-Orbea’s group [26]. The cell lines were maintained in culture at 37°C in a humidified incubator with 5% CO2. Cells were cultured in complete IMDM-glutamax medium with all buffered conditions as reported previously. These cells show resemblance in expression of surface markers and mimic splenic ex-vivo immature CD8α^+^ DCs as shown by extensive characterization done by Prof. Hans Acha-Orbea’s group.

B16F10 cell line obtained from Dr. Shantibhushan Senapati were maintained in DMEM media and were cultured and maintained at 37°C in a humidified incubator with 5% CO2. For *in vitro* experiments, the DCs were plated in 12- or 6-well plates at a density of 5×10^5^ or 1×10^6^ cells/ml overnight. The cells were then challenged with different activation media containing TLR9 agonist CpG-B at a concentration of 1ug/ml, TLR3 agonist pIC at 2ug/ml for 2, 6 or 12 h. For performing RT-qPCR analysis the cells were washed in the plate once with PBS followed by addition of RNA-later (LBP) lysis buffer for lysis of cells. The plates were then stored at −80°C until further RNA isolation and processing of samples.

### Generation of Stable SMRT KD CD8α^+^ MutuDCs

For generating stable SMRT knockdown and their comparative control DC, lentiviral vector pLKO.1 (Sigma) containing three different sigma mission shRNA for *Ncor2* were picked targeting chromosome 5 on mouse genome against exons 48, 19, and 14 respectively (Key Resources Table). Viral particles packaged with shRNA expressing transfer plasmids were produced in 293T cells using Cal-Phos (CaPO4) mammalian transfection kit according to an optimized protocol. We used a 2^nd^ generation lentiviral system which included PCMVR and PMD2G as packaging and envelope plasmids respectively. Human embryonic kidney (HEK) 293T cells were transfected with transfer plasmids containing three different *Ncor2* shRNAs or control shRNAs along with pCMVR8.74 and pMD2G. After 12–14 h the culture medium was replenished and supernatant containing viral particles were collected after 24 h in 50 ml conical tubes. Viral particle-containing culture supernatant was concentrated using ultracentrifugation at 50,000g at 16°C for 2h and preserved at −80°C in small aliquots. For transduction of shRNA containing viruses in CD8α^+^ cDC1 MutuDC lines, the cells were plated at a density of 1.5 × 10^5^ cells/well of 12 well plate followed by transduction with virus particles containing supernatant. The media was replaced with fresh media after 12h of virus incubation with DCs followed by addition of 1 μg/ml puromycin selection medium after 72 h of media replacement for stable KD cells.

### RNA Isolation and RT-qPCR

The extraction of RNA was done using NucleoSpin RNA Plus miniprep kit (Machery Nagel). Briefly, cells were preserved in LBP lysis buffer in −80°C and thawed by placing the plates/tubes on ice. Total RNA was isolated according to the manufacturer’s protocol. RNA concentration was estimated by nanodrop (Thermo) and then 1-2 μg of total RNA was used to prepare cDNA using high-capacity cDNA Reverse Transcriptase kit (Applied Biosystems). Quantitative PCR was performed using SYBR Green master (Roche) and PCR amplification was monitored in real-time using LightCycler-480 Instrument. Primer oligonucleotides for qPCR were designed using the universal probe library assay design system and the primer pairs used are listed in Key Resources Table. Primers were optimized for linear and single product amplification by performing standard curve assays.

### Flow Cytometry (FACS)

We performed flow cytometry analysis using the well-established surface and intracellular (IC) staining protocols [25]. 5 x 10^5^ and 1.5 x 10^6^ cells were seeded for surface and IC staining respectively. Cells were either left unstimulated or stimulated with CpG or pIC for 6h. For staining the cells were dissociated and washed with FACS buffer (3% FCS in 1X PBS, 5 mM EDTA). After washing, fluorochrome conjugated antibodies for proteins of interest were added to the cells as a cocktail in the staining buffer. For surface staining cells were stained in FACS buffer for 30 min in dark at 4°C. For IC staining of cytokines the cells were first fixed with 2% paraformaldehyde for 20 min followed by permeabilization using 1x permeabilization buffer (eBiosciences). The fixed and permeabilized cells were then resuspended in IC staining buffer and stained with fluorochrome tagged antibodies for selected cytokines. For optimal staining the cells were incubated with antibodies for 30 min in dark. After incubation the cells were washed twice with FACS wash buffer and then acquired for differential expression analysis using LSRII fortessa flow cytometer (BD Biosciences). The acquired data was analyzed using FlowJo-X software (Treestar). Antibodies used for flow cytometry experiments are listed in the Key Resources Table.

### Bio-Plex Assay for Cytokine Quantitation from Cell Culture Supernatants

Bio-Plex assay (multiplex ELISA) was used to estimate the cytokine levels secreted in the cell culture supernatants of SMRT KD and control DC and BMcDC1 after 6 h of CpG stimulation according to previous reports [25]. After culture, the supernatants were stored at −80°C in small aliquots until analysis. Cytokine levels were estimated using 23-plex-mouse cytokine assay kit following the vendor recommended protocol (Biorad).

### Generation of Bone Marrow Derived DCs (BMDCs) for *ex vivo* Studies

Six to eight-week-old female C57BL/6 mice were killed by cervical dislocation and disinfected using 75% ethanol [25]. In short the tibias and femurs were removed under sterile conditions, then soaked in RPMI-1640 medium supplemented with 10% FBS. Cells from both ends of the bone were flushed out with a needle of 1-mL syringe from the bone cavity into a sterile culture dish with RPMI-1640 medium. The cell suspension in the dish was collected and centrifuged at 350g for 5 min, and the supernatant was discarded. The cell pellet was suspended with a 1x RBC lysis buffer (Tonbo) for 5-10 min on ice. Cell clumps were then passed through a 70μm strainer to obtain single cell suspensions. The lysed cells were washed once with RPMI-1640, counted and used for differentiation into DCs.

We followed a well-established protocol for differentiation of BMDCs with slight modifications. The cells, suspended in RPMI-1640 medium supplemented with 10% FBS, were distributed into 6-well plates at a density of 1 × 10^6^ cell/ml/well. Subsequently, 1μl/ml of FLT3L containing sera was added into the medium. The cells were cultured at 37°C in an incubator containing 5% CO2 and left untouched for 5 days. On day 5, the suspended and loosely attached cells were collected.

The cells were plated into 96-well plate for lentiviral transduction using concentrated viruses at a density of 0.4 x 10^6^ cells/well for each *Ncor2* shRNA and control shRNA. After 72h the cells were stimulated with CpG for 6h and then immune-profiling was performed using flow cytometry.

### Co-culture of DCs with CD4^+^ T-Cells and CD8^+^ T-cells for Assessing T-Cell Proliferation and Differentiation

DC-T-cell co-culture experiments were performed according to well established protocol [56, 57]. Naïve CD4^+^ or CD8^+^ T-cells were purified from spleen of TCR-transgenic OT-II or OT-I mice using CD4^+^ or CD8^+^ T-cell isolation kit. SMRT KD and control CD8α^+^ cDC1 DCs were seeded at a density of 10,000 cells/well in round bottom 96 well plates followed by pulsing with OVA peptide (323-339) /OT-II at 200nM concentration or OVA peptide (257–264) /OT-I was used at 5nM overnight. Further DCs were stimulated with CpG or pIC for 2h. After 2h, purified OT-II or OT-I T-cells were added at the density of 100,000 cells/well (1:10 ratio). Then T-cell proliferation and differentiation into distinct Th subtypes Th1, Th2, Th17 and Tregs, in case of OT-II, and cytotoxic T-cells, in case of OT-I, were analyzed by FACS. Proliferation was measured using an amine based dye (eFluor 670). The rate of T-cell proliferation was inversely proportional to the median fluorescence intensity (MFI) measured in FACS after 72h of co-culture. For Th and cytotoxic T-cell differentiation profiling after 96h, the co-cultured T-cells were re-stimulated with PMA (10 ng/mL) and ionomycin (500 ng/mL) and followed by Brefeldin-A (10μg/mL) treatment for 5h to block the IC cytokines from being secreted. After 5h, fluorochrome conjugated antibodies specific to different T-cell subtypes were used to profile T-cells into Th1 (T-bet and IFN-γ), Th2 (GATA3, IL-13), Tregs (CD25, FoxP3, IL-10) and Th17 (RORγT, IL-17) or cytotoxic T-cells (perforin, IFN-γ, Granzyme-B). For gating effector T-cells we used CD44 as a marker.

### Chromatin Immuno-Precipitation (ChIP) for p-STAT3

The ChIP for p-STAT3 was performed according to the methods optimized previously by Raghav and Meyer’s lab. For ChIP assays, 40 x 10^6^ CD8a+ cDC1 MutuDCs were seeded in 15 cm^2^ plates and prepared for four ChIP assays by 10 min cross-linking with 1% formaldehyde (sigma) at room temperature followed by quenching using 2.5 M glycine (sigma) for 10 min. The plates were placed on ice and the cells were scraped and collected in 50 ml conical tubes. The cells were then washed three times using cold 1x PBS at 2,000 rpm for 10 min at 4°C and the cell pellets were stored at −80°C. At the day of the ChIP experiment, the cells were thawed on ice followed by lysis using Farham lysis buffer (5 mM PIPES pH 8.0, 85mM KCl, 0.5% NP-40 supplemented with protease and phosphatase inhibitors (Roche)) made in miliQ. The supernatant was aspirated and the pellet was resuspended in RIPA buffer (1% NP-40, 0.5% sodium deoxycholate, 0.1% SDS supplemented with Roche protease and phosphatase inhibitor tablet just before use). The chromatin was fragmented using a Bioruptor (Diagenode) sonicator for 30 min using high amplitude and 30s ON & 30s OFF cycles to obtain 200-500 bp size fragments. A cooling unit was used to circulate the cold water during sonication to avoid decrosslinking because of overheating. After sonication, chromatin length was checked in agarose gel. The fragmented chromatin was centrifuged at 10,000 rpm for 5 min and then clear supernatant was collected in 15 ml conical tubes. The DNA concentration of the chromatin was estimated using a Nano-Drop (Thermo) and the chromatin was diluted with a RIPA buffer to use 150 μg/ml of chromatin for each IP. M-280 sheep anti-rabbit IgG dynabeads 40ul/IP was taken in a 1.5ml MCT tube. 1ml RIPA buffer was added to the beads and placed on a magnetic stand. The MCTs were inverted 5 times and allowed to stand for 3 min. The beads were washed in the same way 3 times. After the 3rd wash the beads were centrifuged shortly and the remaining RIPA buffer was aspirated. To the beads, 5μl of mouse monoclonal anti-p-STAT3 (CST) was added to immunoprecipitated the chromatin complex at 4°C for 8h on rocker shaker. After 8h incubation, the beads were again placed on a magnetic stand and washed with RIPA to get rid of the unbound antibody. Chromatin was added to the beads and placed on a rotating rocker at 4°C overnight. Next day the tubes containing chromatin, antibody, and beads were taken out, placed on a magnetic stand and supernatant was aspirated. The beads were washed 5 times with LiCL IP wash buffer (100mM Tris pH7.5, 500mM LiCl, 1% NP-40, 1% sodium deoxycholate in miliQ) and 2 times with TE buffer (10mM Tris pH7.5, 0.1mM EDTA pH8 in miliQ). After removing the wash buffer completely, protein-bound chromatin complexes were eluted from beads using an elution buffer (1% SDS, 0.1M NaHCO3 in milli-Q water). The chromatin was incubated at room temperature for 30 min in an elution buffer. A short spin was given and the MCT was again placed on a magnetic stand to collect the eluted chromatin. The eluted chromatin was then reverse crosslinked by incubating the eluted supernatant at 65°C overnight on a heat block after adding 8 μl of 5 M NaCl. Next day DNA was purified from the reverse cross-linked chromatin by proteinase-K and RNase digestion followed by purification using PCR purification kit (Qiagen). The purified DNA was eluted in 40μl of elution buffer.

### Chromatin Immuno-Precipitation (ChIP) for SMRT

The ChIP for SMRT was performed according to the methods optimized previously by Raghav and Deplancke’s lab [25, 58]. In short the cells were lysed in a nuclei extraction buffer for 10 min at 4°C while shaking to isolate the nuclei. The isolated nuclei were then washed using a protein extraction buffer at room temperature for 10 min. Washed nuclei were resuspended in chromatin extraction and incubated for 20 min on ice. The chromatin was fragmented using a Bioruptor (Diagenode) sonicator to obtain 200-500 bp-sized fragments. The fragmented chromatin was centrifuged at 17,000g for 10 min and then clear supernatant was collected in chilled 15ml falcon tubes. The DNA concentration of the chromatin was estimated using a NanoDrop and the sonicated chromatin was diluted with ChIP dilution buffer to get 100 μg/ml of chromatin for each IP. BSA and ssDNA (Salmon Sperm DNA) -preblocked protein-A sepharose (80 μl/IP) beads were added to the samples and incubated for 2h to remove non-specific-binding chromatin. To the supernatant, 5 μl/IP rabbit polyclonal anti-SMRT antibody (Abcam) was added to immuno-precipitate the chromatin complex at 4°C overnight. After the overnight incubation, 50μl blocked beads were added to each sample and incubated for 90 min at 4°C to pull down the respective antibody-chromatin complexes. The beads were then washed four times with a low salt wash buffer followed by two washes with high salt wash buffer, lithium chloride wash buffer and tris-EDTA (TE) buffer. After removing the wash buffer completely, protein-bound chromatin complexes were eluted from beads for 30 min using an elution buffer. The eluted chromatin was then reverse-crosslinked by incubating the eluted supernatant at 65°C overnight on a heat block after adding 8μl of 5M NaCl. The next day, DNA was purified from the reverse crosslinked chromatin by proteinase and RNase digestion followed by purification using Qiagen DNA purification columns. The purified DNA was eluted in 50μl of Qiagen elution buffer.

### ChIP/RNA-seq library preparation for sequencing

For RNA-seq library preparation 2ug of total RNA was used to isolate mRNA using magnetic beads with mRNA isolation kit (PolyA mRNA isolation module, NEB). Later mRNA library preparation kit, NEB, was used for RNA-seq library preparation according to manufacturer’s protocol. Concentration of the libraries were estimated by Qubit 2.0 (Invitrogen) and fragment sizes were analysed in Bio-analyzer (Agilent). The libraries were then sequenced on Illumina NextSeq 550 platform.

Similarly for ChIP seq 30 μl ChIP-DNA was processed for library preparation according to ChIP-seq library preparation protocol (NEB) [25]. After library preparation and quality check, the libraries were sent to NGS service provider (Sci Genome, Bangalore, India) for Illumina sequencing using NextSeq-550 instrument.

### Western Blotting

Cells were collected in RIPA buffer (0.5 M EDTA, 1 M Tris-Cl pH7.5, 1 M NaCl, 200 mM, Roche protease inhibitor) at 0h, 2h and 6h CpG stimulation. Cells were lysed completely by sonication of the samples in Bioruptor (Diagenode) for 10 min using high amplitude and 30s ON & 30s OFF cycles. Protein concentrations were measured in 96 well plates using BCA protein assay kit (BioRad) at 562nM. For western blot of phospho and its respective total protein molecule we first probed the membrane with phosphoantibodies, stripped and re-probed the same membrane with respective total antibodies. For densitometric analysis we first normalized phosphorylated form of STAT3 /mTOR with their respective loading controls. The similar approach was followed for its corresponding total protein. Finally the ratio of normalized values were plotted as relative intensity..

### Delayed Type Hypersensitivity (DTH) Assay

DTH was performed using culture grade ovalbumin (OVA) from chicken egg (Sigma) dissolved in 1x PBS at a concentration of 1mg/ml and filtered through 0.2-micron PES syringe filter. 1.5ml alum and 1.5ml OVA was added in a glass beaker and passed through a glass syringe multiple times to make an emulsion. 300μl per mice was injected subcutaneously in the back behind ears in each mice for OVA immunisation. After 14 days control and SMRT KD DCs were pulsed with OVA (100ug/ml) for 4 h. Cells were then stimulated with CpG. After 2h of stimulation the cells were dissociated and injected at 10 x 10^6^ cells/mice. Further after 7 days OVA (20mg/ml) was heated at 80°C for 2h, cooled, and injected in foot pad of mice (25μl/mice). 1x PBS was injected in the alternative footpad. Paw thickness was measured till 72h using Vernier caliper. After 72h the popliteal and inguinal lymph nodes were isolated and checked for T-bet IFN-γ as well as RORγt IL-17.

### OVA specific ELISA

To examine OVA specific immune response we performed experiments as described [25, 59]. In brief, we collected sera at day 20 and day 23 after OVA immunization to perform ELISA for OVA specific IgG titer. Elisa plates were coated with 100ug/ml of OVA (Sigma) prepared in a coating buffer (Na_2_CO_3_, NaHCO_3_, Sodium Azide) overnight at 4°C following five washes with washing buffer (PBS with 0.05% tween −20). Blocking was done with PBST containing 0.5% gelatin for 1h at 37°C. After five times washing, 50μl diluted sera were added from mice and kept for 1.5 h at 37°C. IgG1 (dilution 1:10,000) and IgG2a (dilution 1:100) was detected using biotin labelled anti-mouse IgG1 and IgG2a while total IgG (dilution 1:1000) was detected using anti-mouse HRP conjugated IgG followed by anti-mouse streptavidin-HRP (Biolegend). The plates were further incubated at 37°C for 1h and washed 7 times with a washing buffer. Color was developed by TMB (50μl/well) and incubated in dark for 10 min. 2N H_2_SO_4_ (50μl/well) was used to stop the reaction. The plates were read using ELISA reader for IgG estimation at 450nm.

### Tumor cell lysate preparation

Tumor lysate was prepared from previous reports with some modifications [60]. B16/F10 tumor cells were adjusted to 3□×□ 10^6^ cells/ml in DMEM medium. Cells were subjected to 3 freeze (-□80°C) / thaw (40°C) cycles of minimum 20 min each. The lysed cells were checked under trypan blue staining and centrifuged at 12,000rpm for 15 min. The supernatant was passed through a 40μm cell strainer before adding to cDC1s seeded at a density of 3 x 10^6^ (DC:tumor cell ratio of 1:1).

### B16F10 Tumor Model

Mice were injected subcutaneously (s.c.) with 0.5 x 10^6^ tumor cells into the left flank, and a booster dose was re-injected 3 days later. 7 days after booster dose, 0.1 x 10^6^ cells were injected subcutaneously in mice in the right flank. Tumor growth was measured every day using a vernier caliper till 16 days. Tumors were removed, weighed, and dissociated to make single cell suspensions.

### Isolation of tumor and tumor re-stimulation

For IC perforin, granzyme-B, and IFN-γ staining, single cell suspensions were *ex vivo* restimulated with 10ng/ml PMA, 500ng/ml ionomycin, and 5□μg/mL Brefeldin A for 5h. Cells were labelled with indicated surface-staining antibodies, fixed with 2% PFA, permeabilized with permeabilization buffer and stained with IC antibodies.

### RNA-seq data processing and analysis

Raw reads of SMRT KD RNA-seq samples and its matched control in unstimulated and 6h CpG stimulation were checked for quality using FASTQC [61], and aligned to mouse genome (UCSC mm10) using hisat2 [62] (with default parameter). Similarly, raw reads of NCoR1 KD and its matched control RNA-seq were processed for quality control and alignment. Raw counts of genes were extracted using featureCount (featureCounts -p -B) [63]. Principal component analysis was performed on variance stabilized transformed (vst) values from DESeq2 [64] using the plotPCA function and plotted using ggplot2 [65]. Further, differential gene expression analysis was performed between NCoR1/SMRT KD compared to its matched control in unstimulated and 6h CpG stimulation condition. Genes were filtered based on log2foldchange (upregulated >= 1 and downregulated <= −1) and adjusted P-value (< 0.05). Total differentially expressed genes were combined from all the comparisons and unsupervised k-means clustering were performed based on log2foldchange values and divided into six clusters. Pathway enrichment analysis for each cluster was performed using Ingenuity pathway analysis.

### ChIP-seq data processing and analysis

SMRT ChIP-seq raw reads in unstimulated and 6h CpG stimulation were checked for quality using FASTQC and aligned to mouse genome (RefSeq mm10) using bowtie2 [66]. Reads were filtered using MarkDuplicates function of Picard and mapping quality >=10 using SAMtools [67, 68]. Peak calling was performed using findPeaks (Homer) and factor as style. Peak calling for NCoR1 ChIP-seq data from our previous study were performed with reads down sampled to the level of SMRT i.e. 9M. Distribution analysis of peaks based on distance relative to TSS were performed using ChIPSeeker [69]. Peaks from both NCoR1 and SMRT were merged using bedops merge command and consensus peaks were generated for further downstream analysis [70]. GeneOverlap R package were used to identify differentially expressed genes that are the direct target of SMRT [71].

### ChIP-seq Peak analysis

Differential binding analysis for NCoR1 and SMRT were carried out on the merged peak using the getDifferentialPeaks program of Homer with cut-off of 2-fold enrichment over background [72]. Peaks were filtered out that didn’t show any differential binding in any of the comparisons. Differential peaks were then categorized based on fold change. Peaks from different categories were annotated to nearest genes using the ChIPseeker R package [69]. De novo motif enrichment analysis was performed using findMotifs.pl (size −50, 50 -len 8, 10, 12) using background generated from provided input genomic regions. P-value <=1e-10 were used to filter significantly enriched TF motifs. KEGG pathway enrichment analysis of differentially expressed genes associated with each binding category were performed using clusterProfiler R package [73].

## Declarations

### Ethics approval and consent to participate

All the animal experiments were performed after getting due approval from the institutional animal ethics committee. All human subjects provided informed consent and the study was approved by the Institutional Ethics Committee.

### Availability of Data and material

RNA-seq and ChIP-seq data of SMRT samples has been submitted to ArrayExpress and available under accession number E-MTAB-10070 (http://www.ebi.ac.uk/arrayexpress/experiments/E-MTAB-10070), E-MTAB-10069 (http://www.ebi.ac.uk/arrayexpress/experiments/E-MTAB-10069), E-MTAB-10864 (http://www.ebi.ac.uk/arrayexpress/experiments/E-MTAB-10864). RNA-seq and ChIP-seq data of NCoR1 samples are available at Gene Expression Omnibus having accession number GSE110423. The R code used in this study has been deposited to https://github.com/sraghav-lab/NCoR1-and-SMRT-Project.

## Acknowledgments

We are sincerely thankful to Hans Acha-Orbea for providing us the DC cell lines, OT-I, OT-II, and FLT-3 transgenic mice. We would like to thank Subhasish Prusty for helping in western experiments and Niyati Das for helping in animal experiments. A.J. is supported by ILS fellowship, A.A. is supported by DBT-SRF, G.P.M. is supported by DBT BINC fellowship, K.S. is supported by UGC-SRF, S.S. is supported by ILS fellowship, M.A. is supported by CSIR funding, V.K.B. is supported by ILS tribal flagship fellowship, A[1].T. is supported by DST INSPIRE.

## Supplementary File legends

**Supplementary File 1:**

Excel file having list of differentially expressed genes (DEGs) in SMRT KD compared to control at 0h and 6h CpG stimulation.

**Supplementary File 2:**

Excel file having a list of annotations of direct target genes that are bound and differentially expressed upon SMRT KD at 0h and 6h CpG stimulation.

**Supplementary File 3:**

Excel file having list of enriched Ingenuity pathway terms for direct target genes of SMRT that upregulated and downregulated in SMRT KD compared to control at 6h CpG stimulation.

**Supplementary File 4:**

Excel file having list of DEGs divided in 6 k-means clusters based on Log2 foldchange in NCoR1 KD and SMRT KD compared to Control DCs in Uns and 6h CpG stimulated condition

**Supplementary File 5:**

Excel file having list of enriched IPA pathway terms for all the 6 k-means clusters.

**Supplementary File 6:**

Excel file having list of enriched KEGG terms for list of direct target genes of differential binding sites of NCoR1 and SMRT.

