## Supplementary figures and images for "NCoR1 and SMRT fine-tune inflammatory versus tolerogenic balance in dendritic cells by differentially regulating STAT3 signaling"

### Supplementary Figure 1

Supplementary Figure 1

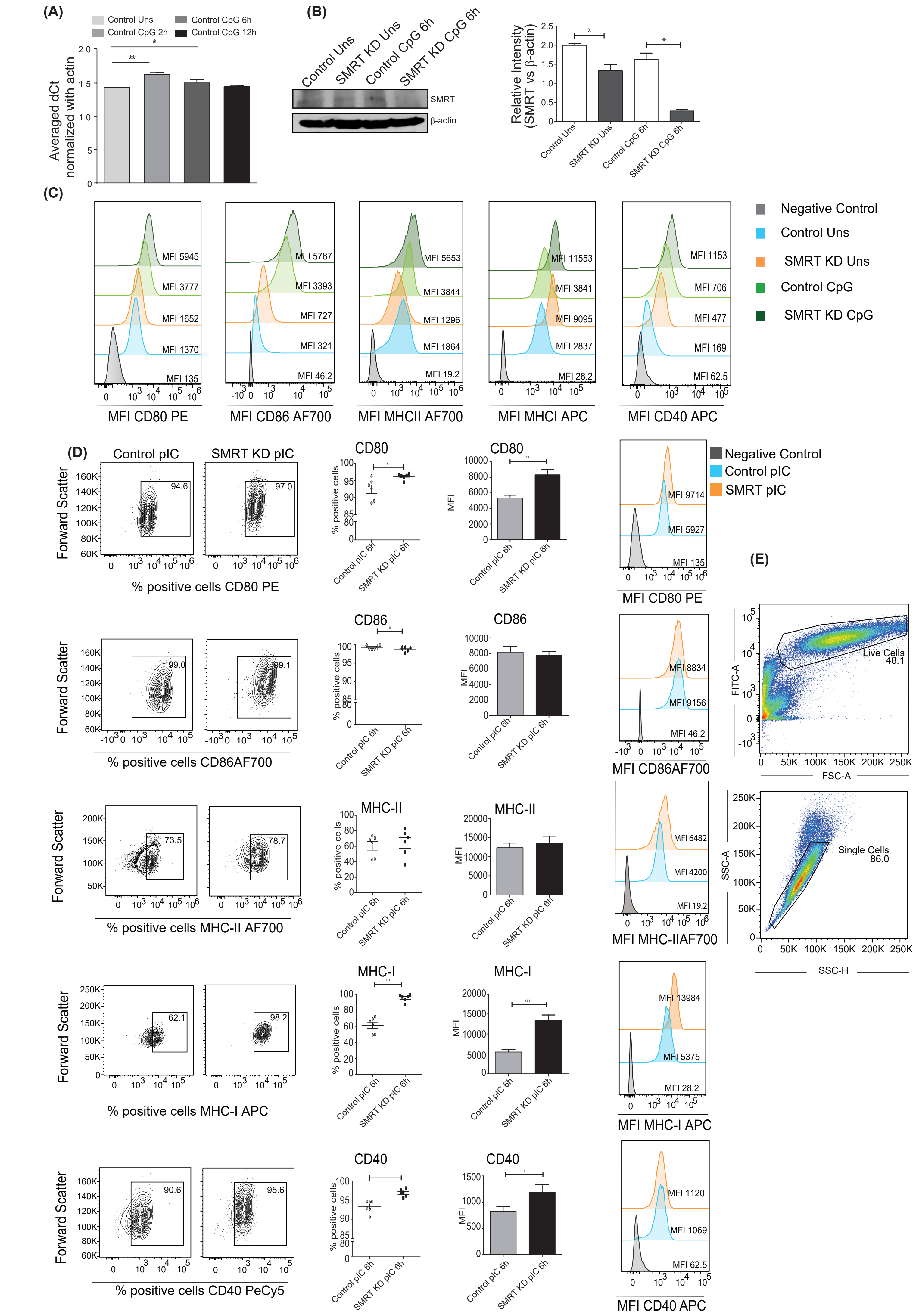
