## Supplementary Figure 2 for "NCoR1 and SMRT fine-tune inflammatory versus tolerogenic balance in dendritic cells by differentially regulating STAT3 signaling"

Figure S1. SMRT depleted cDC1 DCs exhibit enhanced co-stimulation and cytokine secretion upon TLR9 and TLR3 stimulation **(A)** RT-qPCR showing the relative transcript expression of Smrt (averaged dCt normalized with β-actin) in unstimulated and 2h, 6h, and 12h CpG stimulated cDC1s. (n=3) **(B)** Western blot showing SMRT protein expression in unstimulated and 6h CpG stimulated control and SMRT KD cDC1. Densitometry analysis depicted normalized intensity of SMRT bands in SMRT KD cDC1 and control cells. Housekeeping gene β-actin was used as loading control. (n=3) **(C)** Flow cytometry analysis depicting histogram with MFI from flow cytometry analysis of co-stimulatory molecules CD80, CD86, CD40 and activation marker MHC-II, and MHC-I in unstimulated and 6h CpG stimulated control and SMRT KD cDC1s. **(D)** Flow cytometry analysis of co-stimulatory molecules CD80, CD86, and CD40 as well as activation markers MHC-II and MHC-I in 6h pIC stimulated control and SMRT KD cDC1s. Contour plot, dot plot, bar plot, and histogram showing cell population, percent positive cells and MFI respectively. (n=3-6) **(E)** Back gating strategy used throughout the samples.

*p ≤ 0.05, **p ≤ 0.01 and ***p ≤0.001. p-value has been calculated using two tailed paired student’s t-test. Data shown in figure is combined from 2-3 independent experiments [A & C]. Error bars represent SEM.

Supplementary Figure 2

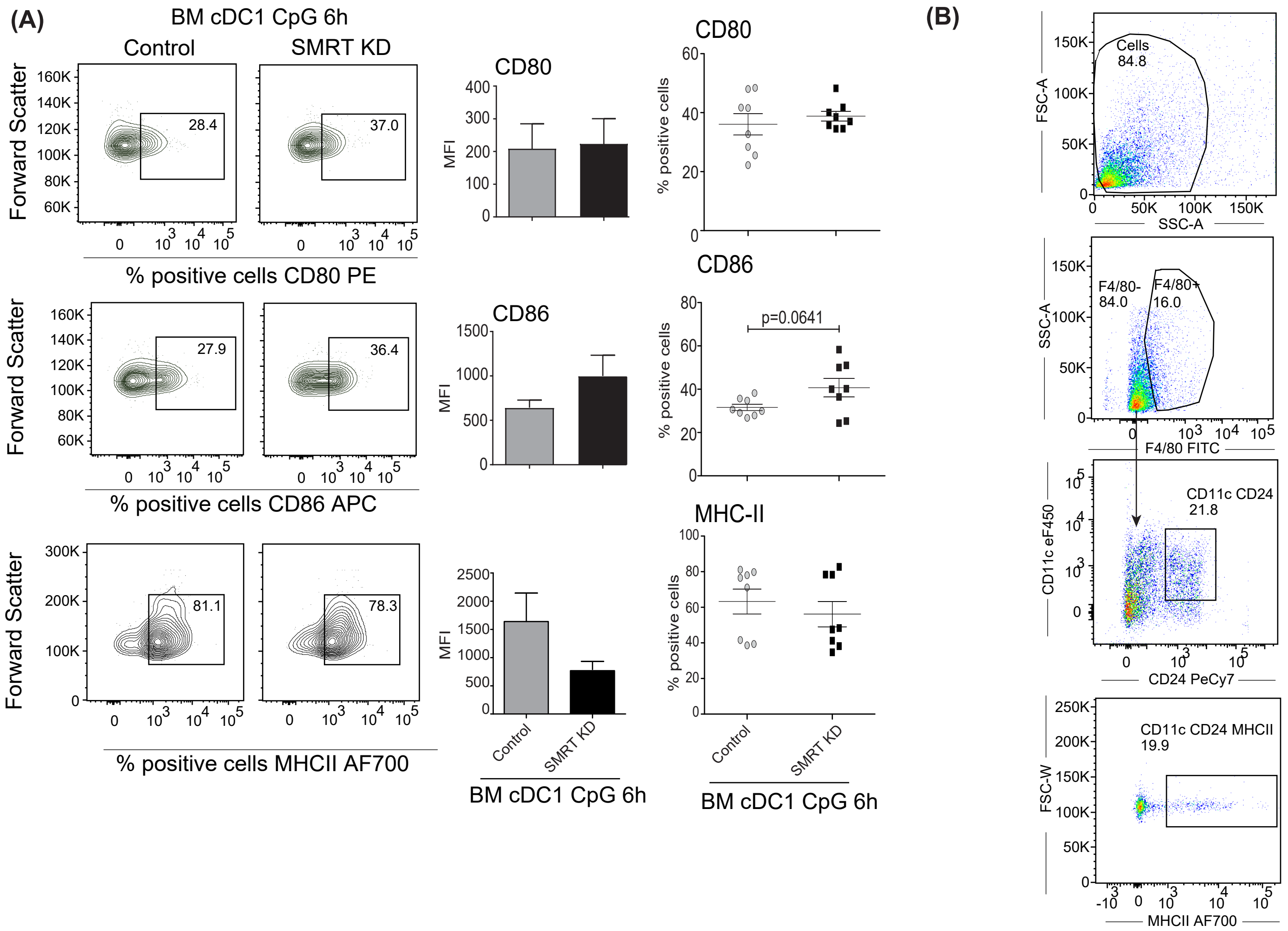


Figure S2. SMRT depleted bone-marrow derived DCs differentiated with FLT3 ligand exhibit enhanced co-stimulation upon CpG stimulation. **(A)** Flow cytometry analysis of activation and co-stimulatory molecules CD80, CD86, and MHC-II in 6h CpG stimulated control and SMRT KD BMcDC1s. Contour plots, bar plots, and scatter dot plots depicting percent positive cells and MFI. (n=8) **(B)** Back gating strategy used throughout BMcDC1 samples.

*p ≤ 0.05, **p ≤ 0.01 and ***p ≤0.001. p-value has been calculated using two tailed paired student’s t-test. Datashown in figure is combined from 4 independent experiments [A]. Error bars represent SEM.
