## Supplementary Figure 3 for "NCoR1 and SMRT fine-tune inflammatory versus tolerogenic balance in dendritic cells by differentially regulating STAT3 signaling"

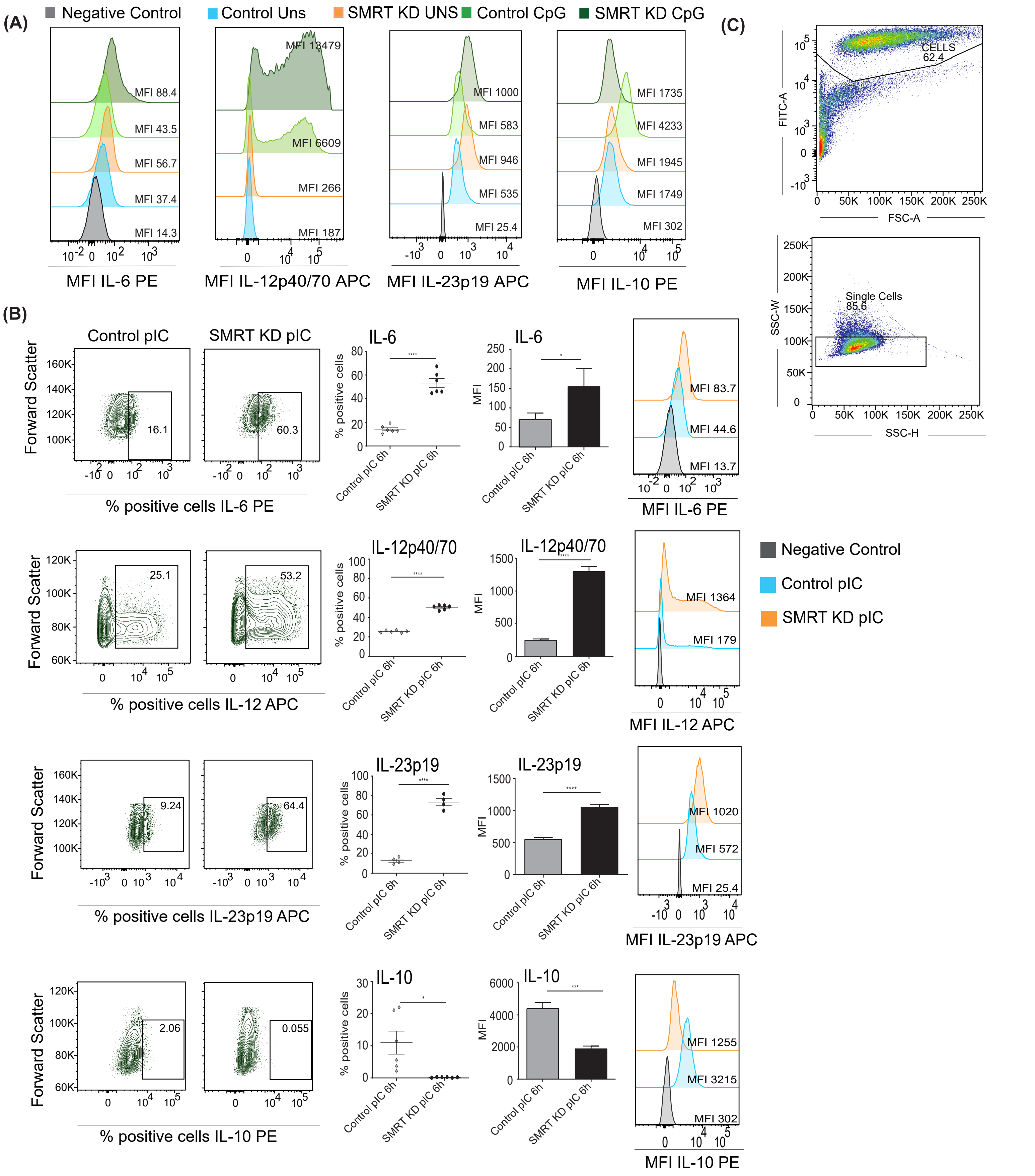


Figure S3. SMRT depleted cDC1 DCs cytokine secretion upon TLR stimulation **(A)** Flow cytometry analysis showing histograms depicting MFI of pro-inflammatory cytokine IL-6, IL-12p40, IL-23p19, and the anti-inflammatory cytokine IL-10 in unstimulated and 6h CpG stimulated control and SMRT KD cDC1s. **(B)** Flow cytometry analysis of pro-inflammatory cytokines IL-6, IL-12p40, IL-23p19, and the anti-inflammatory cytokine IL-10 in 6h pIC stimulated control and SMRT KD cDC1s. Corresponding contour plot, scatter dot plot, bar plot, and histogram showing cell population, percent positive cells and MFI shifts respectively. (n=4-6) **(C)** Back gating strategy used for mutuDC analysis.

*p ≤ 0.05, **p ≤ 0.01 and ***p ≤0.001. p-value has been calculated using two tailed paired student's t-test.
Data shown in figure is combined from 2-3 independent experiments [C]. Error bars represent SEM.
