## Supplementary Figure 4 for "NCoR1 and SMRT fine-tune inflammatory versus tolerogenic balance in dendritic cells by differentially regulating STAT3 signaling"

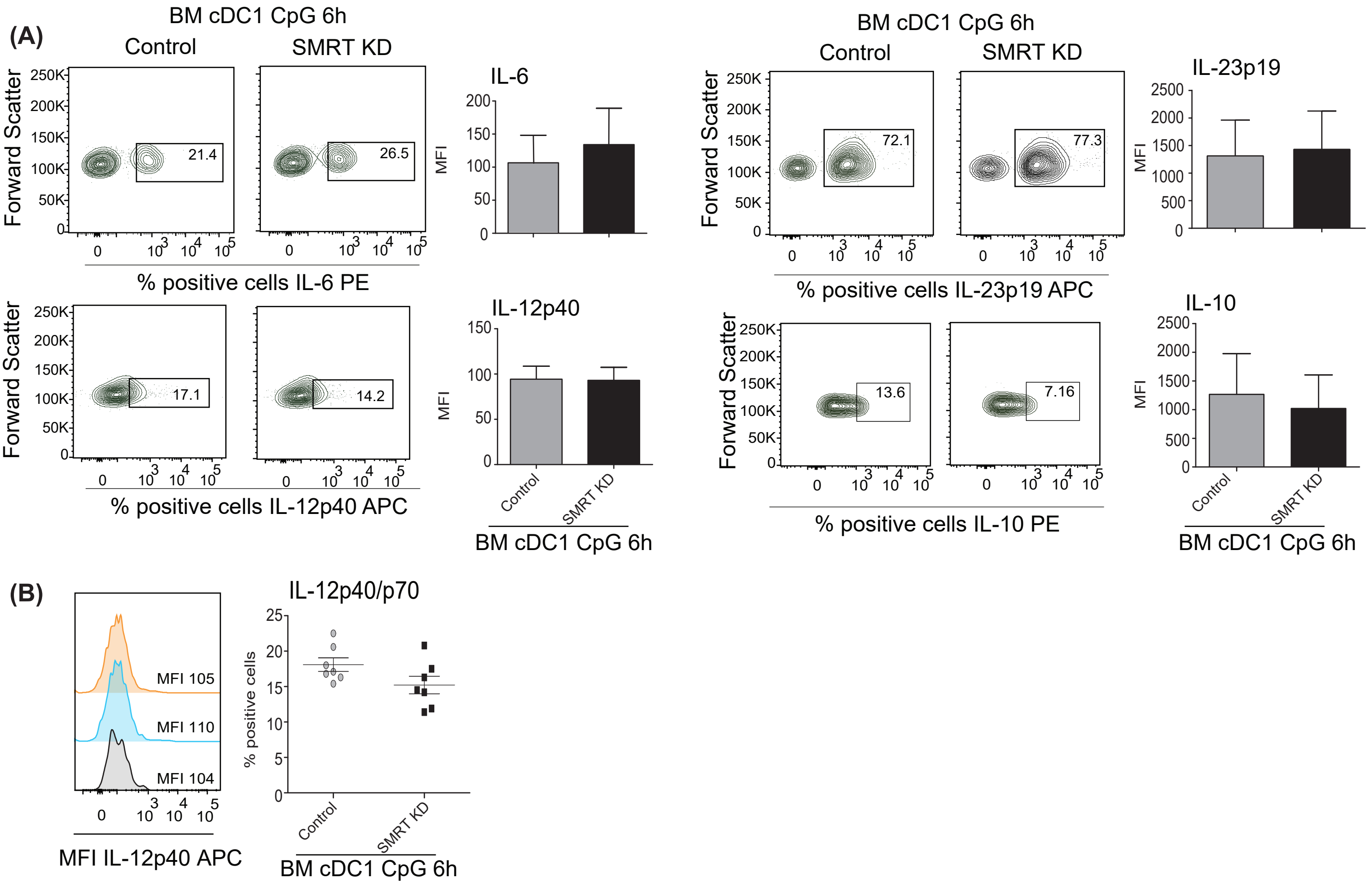


Figure S4. SMRT depleted bone-marrow derived DCs differentiated with FLT3 ligand exhibit enhanced inflammatory cytokine production upon CpG stimulation **(A)** Flow cytometry analysis of proinflammatory cytokine IL-6, IL-12p40, IL-23p19, and the anti-inflammatory cytokine IL-10 in 6h stimulated control and SMRT KD BMcDC1s. Contour plots and bar plot depicting percent positive cells and MFI. (n=3-7) **(B)** Flow cytometry analysis depicting histogram and dot plot of proinflammatory cytokine IL-12p40 in 6h CpG stimulated control and SMRT KD BMcDC1s. (n=7)

*p ≤ 0.05, **p ≤ 0.01 and ***p ≤0.001. p-value has been calculated using two tailed paired student's t-test. Data shown in figure is combined from 2-4 independent experiments [A] and from 4 independent experiments [B]. Error bars represent SEM.
