## Supplementary Figure 5 for "NCoR1 and SMRT fine-tune inflammatory versus tolerogenic balance in dendritic cells by differentially regulating STAT3 signaling"

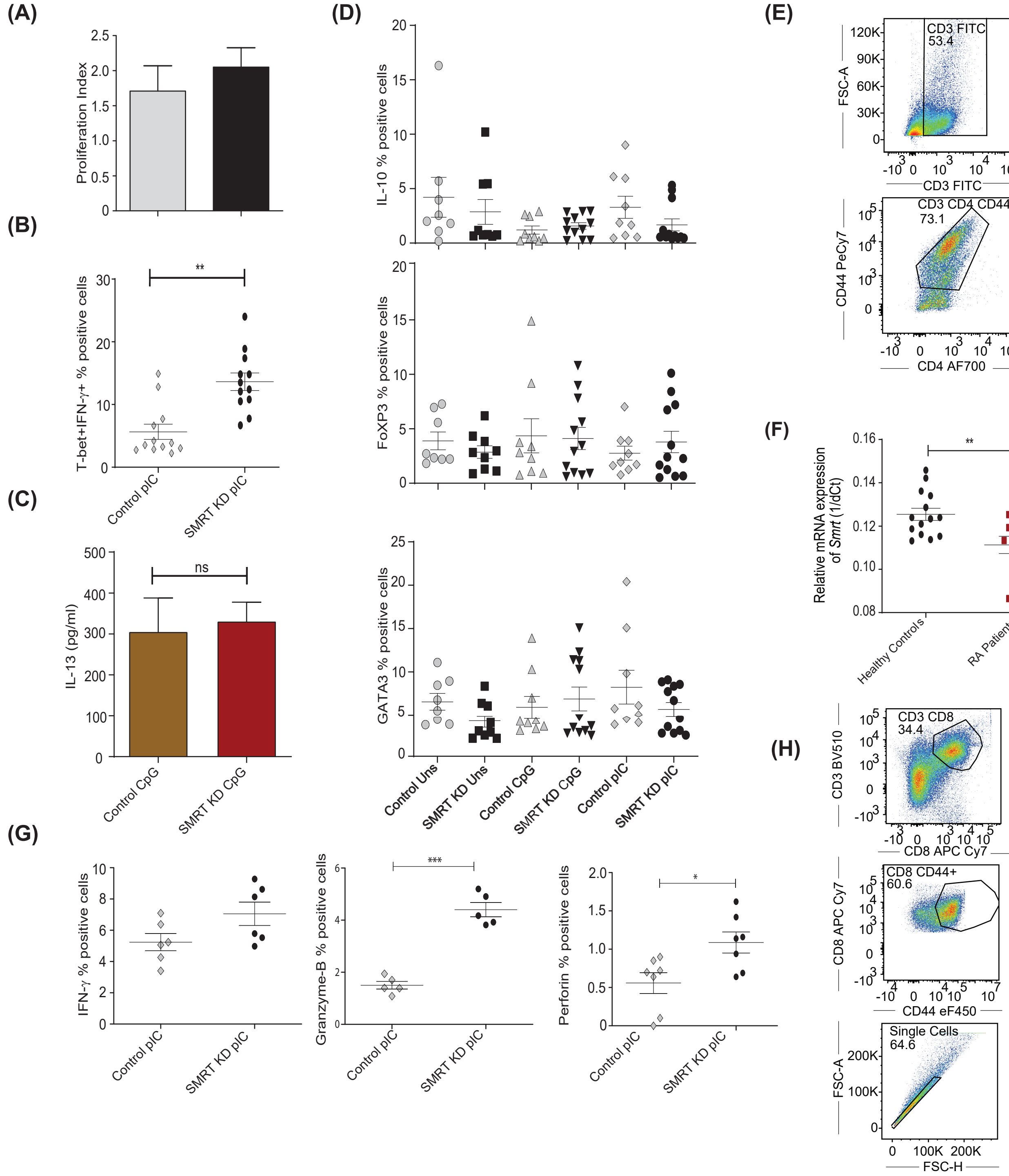


Figure S5. SMRT KD cDC1s increased Th1 and Th17 polarization ex vivo in CD4+ T-lymphocytes and enhanced cytotoxic T-cell activity in CD8+ lymphpcytes. **(A)** Proliferation Index T cells depicting the difference in proliferation rate of OT-II Th-cells co-cultured with control and SMRT KD cDC1s treated with OVA 323-339 peptide overnight followed by pIC stimulation. (n=6) **(B)** Dot plots representing co-cultured OT-II Th-cells showing signature transcription factor and cytokine for Th1, T-bet and IFN-γ response in pIC stimulated condition. (n=12) **(C)** Bioplex cytokine assay depicting quantification of IL-13 cytokine secreted in the culture supernatant of OT-II Th-cells which were co-cultured with control and SMRT KD DCs pulsed with OT-II peptide and challenged with CpG. (n=5) **(D)** Dot plots representing co-cultured OT-II Th-cells showing IL-10, FoxP3, and GATA3 expressing T-cells co-cultured with control and SMRT KD DCs in unstimulated, CpG stimulated, and pIC stimulated condition. (n=8-12). **(E)** Back gating strategy used for flow cytometry analysis of co-cultured OT-II T-cells. **(F)** RT-qPCR showing relative mRNA expression of *Ncor2* transcript (1/dCt) in RA patients (n=11) and their healthy counterparts (n=14). **(G)** Dot plots representing co-cultured OT-I T-cells showing cytokines for cytotoxic T-cells perforin, granzyme-B, and IFN-γ in pIC stimulated condition. (n=5-7) **(G)** Back gating strategy used for flow cytometry analysis of co-cultured OT-I T-cells.

*p ≤ 0.05, **p ≤ 0.01 and ***p ≤0.001. p-value has been calculated using two tailed paired student's t-test.
Data shown in figure is combined from 2 independent experiments [A], from 4 independent replicates [B], from 3 independent replicates [C], from 2 independent replicates [D], and 2 independent replicates [G]. Error bars represent
