## Supplementary Figure 6 for "NCoR1 and SMRT fine-tune inflammatory versus tolerogenic balance in dendritic cells by differentially regulating STAT3 signaling"

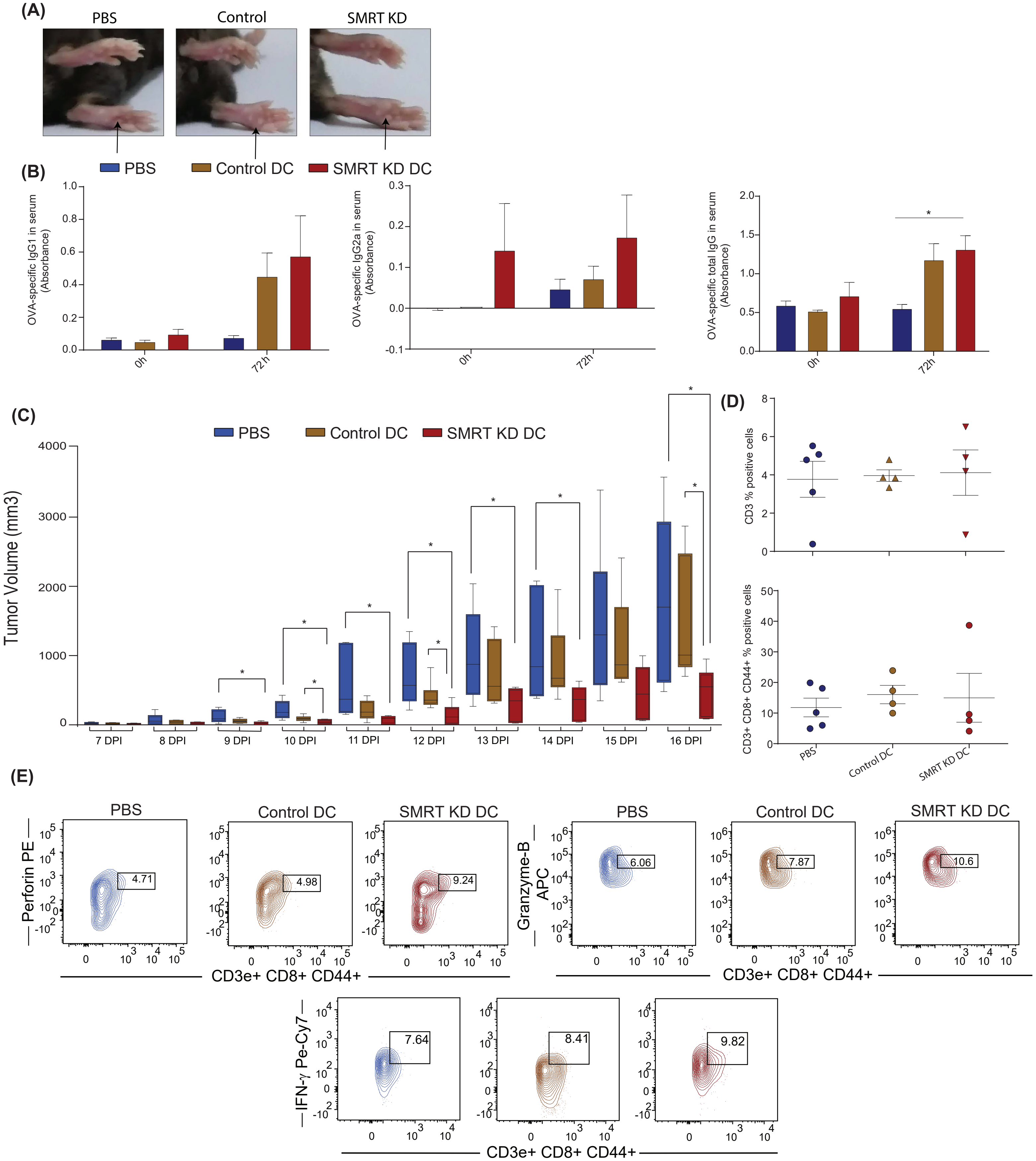


Figure S6. Induction of DTH and B16F10 induced melanoma model in C57BL/6. **(A)** Image showing foot-pad swelling from PBS, ova pulsed control DCs, and SMRT KD DCs treated mice 72h post ova rechallenge. **(B)** ELISA of IgG1, IgG2a, total IgG in 0 and 72h after OVA immunization in in mice injected with PBS, ova pulsed control DCs, and SMRT KD DC. (n= 4-6) **(C)** Box plot showing tumor volume that was taken every day starting from 7th day after B16F10 injection till 16 days post tumor rechallenge in PBS, control and SMRT KD DCs injected mice. **(D)** Dot plots depicting percentage of CD3+ and CD8+ CD44+ cells in all three groups of mice. (n=4-5) **(E)** Contour plots depicting percent positive cells expressing perforin, granzyme-B, and IFN-γ in three groups of mice.

*p ≤ 0.05, **p ≤ 0.01 and ***p ≤0.001. p-value has been calculated using two tailed unpaired student's t-test. Data shown in figure is combined from 2 independent experiments [A-C] and 1 replicates [D]. Error bars represent SEM.
