## Supplementary Figure 7 for "NCoR1 and SMRT fine-tune inflammatory versus tolerogenic balance in dendritic cells by differentially regulating STAT3 signaling"

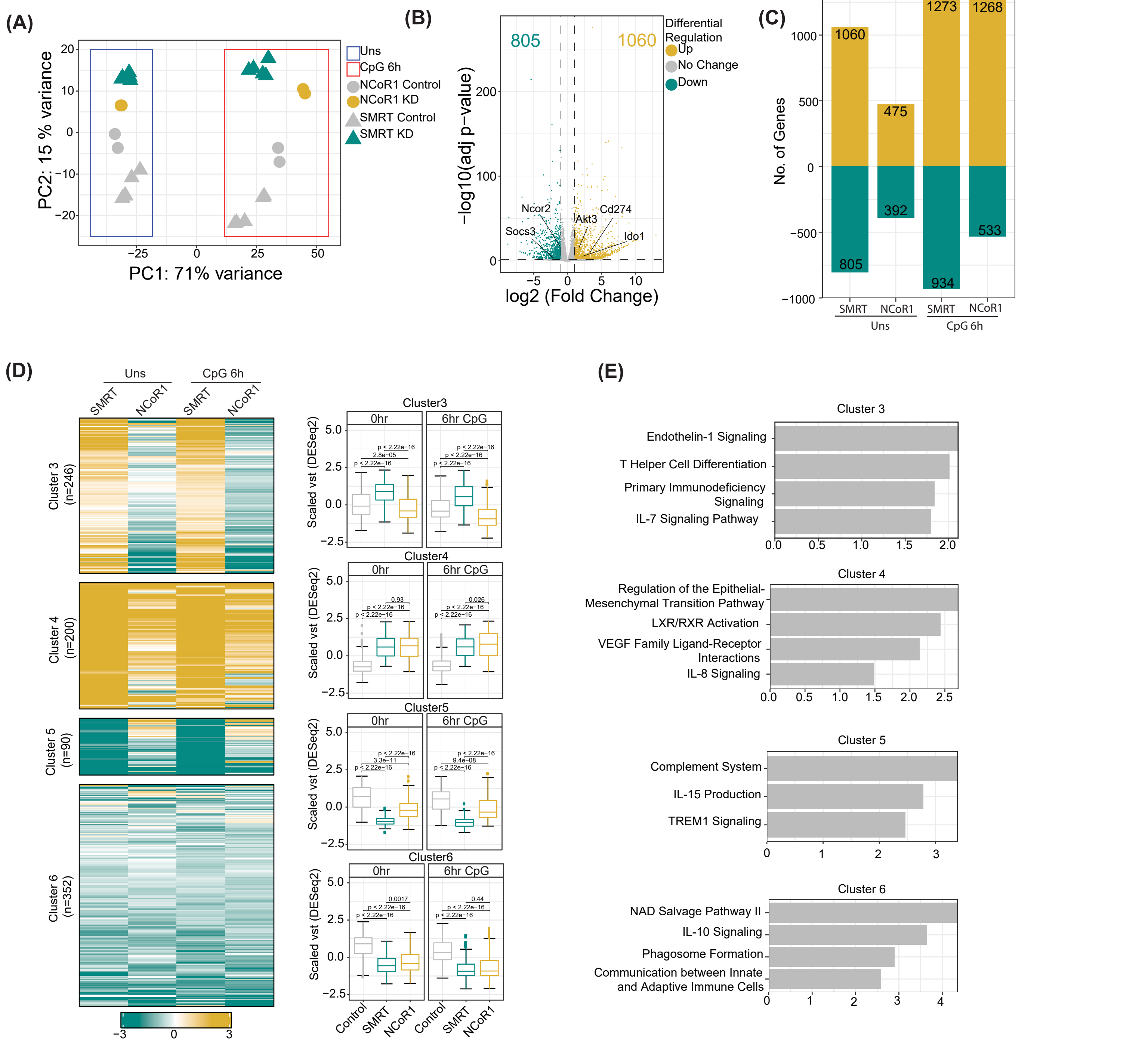


Figure S7. Integrative genomics analysis identified the differential role of SMRT and NCoR1 in regulation of immune response in cDC1. **(A)** PCA plot showing the percent of the variation in principal component 1 and 2 for control, NCoR1 KD and SMRT KD sample in unstimulated and 6h CpG stimulated DCs. **(B)** Volcano plot showing the differentially expressed genes (DEGs) in SMRT KD cDC1 DCs as compared to control cells in unstimulated condition. 1060 and 805 genes were upregulated and downregulated respectively upon SMRT depletion. **(C)** Bar plot showing comparison of number of total DEGs after SMRT KD and NCoR1 KD in unstimulated and 6h CpG stimulation condition. **(D)** Heatmap showing clusters 3-6 of K-means clustering (shown in Fig 6F) of log2 fold change of DEGs upon SMRT and NCoR1 KD compared to control cells in unstimulated and 6h CpG stimulated DCs. The box plot shows scaled normalized expression values (rlog) from DESeq2 for the respective cluster. **(E)** The bar plot showing enriched pathway terms from Ingenuity pathway analysis for respective clusters.
