## Supplementary Figure 8 for "NCoR1 and SMRT fine-tune inflammatory versus tolerogenic balance in dendritic cells by differentially regulating STAT3 signaling"

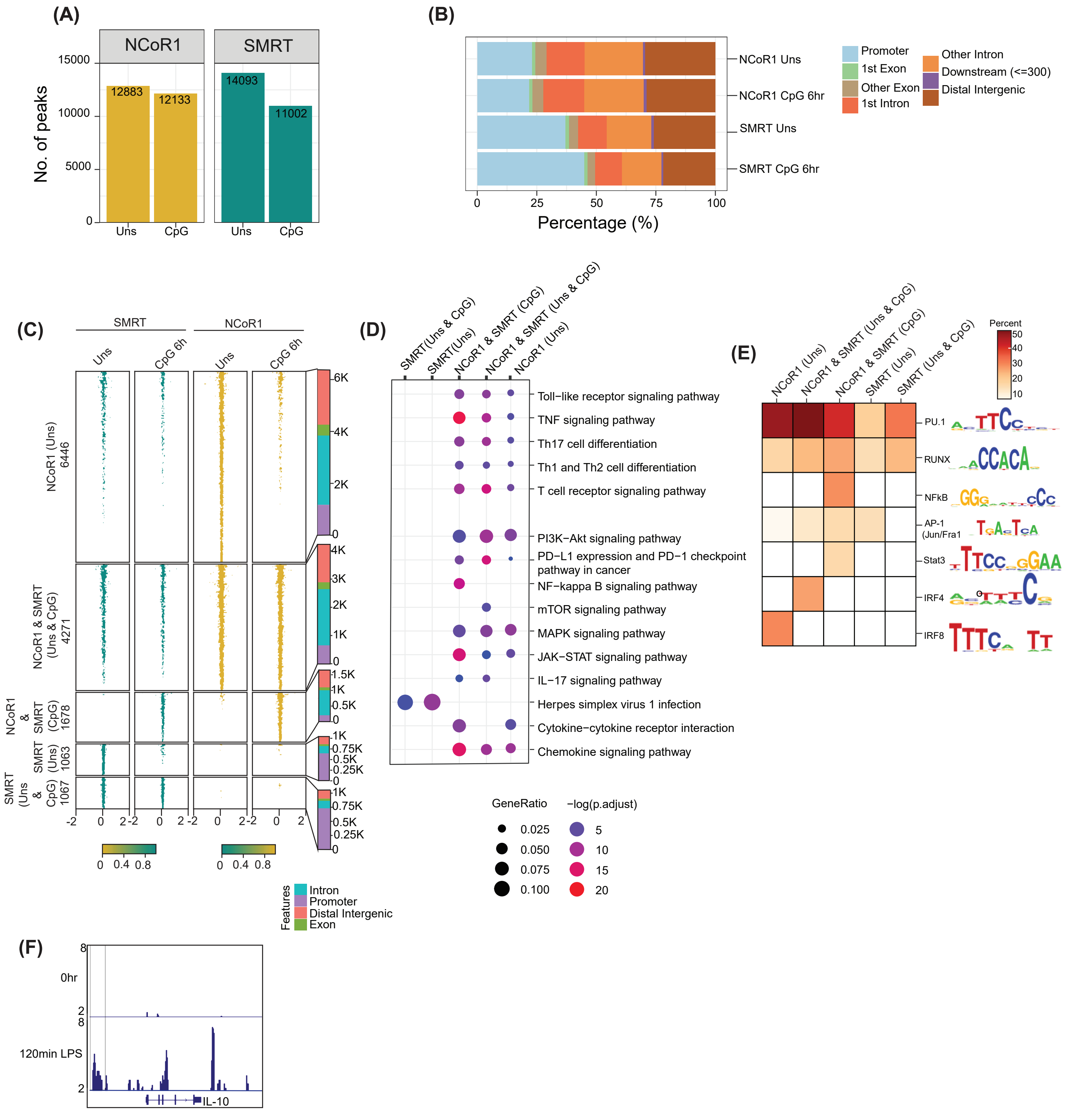


Figure S8. Integrative genomics analysis identified the differential role of SMRT and NCoR1 in regulation of immune response in cDC1. **(A)** Bar plot showing the total number of identified peaks (binding sites) of NCoR1 and SMRT in unstimulated and 6h CpG stimulated DCs. **(B)** Percentage stacked bar plot showing the global distribution of NCoR1 and SMRT identified peaks based on distance relative to TSS in unstimulated and 6h CpG stimulation. **(C)** Tornado plot showing ChIP-seq signal (± 2kb to peak center) of differential NCoR1 and SMRT binding sites in unstimulated and 6h CpG stimulation. Bar plot showing the distribution of differential genomic regions based on distance relative to TSS. **(D)** Dot plot showing the enriched KEGG terms for genes associated with differential NCoR1 and SMRT binding clusters (genomic regions). **(E)** Heatmap showing percent of target binding sites with transcription factor motifs that were significantly enriched (P-value < 1e-10) in differential NCoR1 and SMRT genomic regions. **(J)** IGV snapshot showing ChIP-seq enrichment of STAT3 on Il-10 gene in BMDCs at 0h and 120 min post LPS stimulation.
